# Copper-transporting ATPase ATP7B and the lysosomal exocytosis pathway synergise to detoxify cadmium from cells

**DOI:** 10.64898/2026.05.16.725294

**Authors:** Debosmita Bhattacharya, Kaustav Chakraborty, Raviranjan Pandey, Brindaban Maji, Ashima Bhattacharjee, Arnab Gupta

## Abstract

Cadmium, being a highly toxic metal, perturbs cellular homeostasis by forming stable complexes with numerous thiol-active proteins, ultimately leading to severe liver and lung damage. Despite its well-documented toxicity, the molecular mechanisms governing cadmium export remain poorly understood. Given the chemical similarity between cadmium and copper, we investigated whether the canonical copper-exporting ATPases, ATP7A and ATP7B participate in cadmium handling. Upon Cd treatment in hepatocytes, ATP7B undergoes trafficking to lysosomes via the retromer complex, as also observed in the case of elevated copper, accompanied by the upregulation of acidic lysosomal populations. In contrast, ATP7A expressed in lung adenocarcinoma cells, though exhibit vesicular redistribution upon Cd exposure, does not mediate lysosomal sequestration, suggesting distinct deployment of late secretory pathways by the two copper ATPases in response to cadmium. We have also observed that ATP7B^−/−^ hepatocytes exhibit increased sensitivity to Cd exposure compared to wild-type cells. Whereas, overexpressing the ATP7B amino-terminal copper-binding domain in bacteria alleviates cadmium-induced stress, indicating its capacity to sequester Cd. *Caenorhabditis elegans* lacking copper-ATPase *cua-1,* displayed increased Cd sensitivity, while mutants (*glo-1*^−/−^), deficient in lysosome-related organelles (LRO), and (*lmp-1^−/−^*), deficient in lysosomal membrane glycoprotein, showed reduced resistance to cadmium toxicity. Treatment of the worm with cadmium increases the abundance of lysosomes marked by elevation in lysosomal biogenesis and functional genes, reinforcing the importance of lysosomal pathways in cadmium detoxification. To summarise, we delineated the non-canonical role of copper ATPases and lysosomes in cadmium-induced cellular toxicity.

## Introduction

Cadmium (Cd), an extremely hazardous heavy metal from Group IIB of the periodic table, is widely found in soil, sediments, air, and water, typically in the 0.1–1 mg kg⁻¹ concentration range^1^. Notably, its cationic form Cd(II) poses a more potent threat than elemental mercury due to its high solubility in water, which facilitates its pervasive distribution within organisms^2^. Cadmium contamination of food is more common in polluted agricultural regions, posing a very serious problem worldwide. Population-based studies conducted in heavily polluted areas have linked cadmium to an increased risk of cancer, cardiovascular, and all-cause mortality^3,4,5,6^. 14 to 17% of cropland is affected by cadmium and other toxic metal pollution globally, and estimate that between 0.9 and 1.4 billion people live in regions of heightened public health and ecological risks^7^. The continued use of cadmium in industry drastically affects the environment, resulting in high human exposure to the element.

In recent years, the biological effects of cadmium (Cd) have been widely investigated, as it belongs to the group of toxic, carcinogenic, and endocrine-disrupting elements^8^. The biological half-life of cadmium (Cd) in the human body is remarkably long, ranging from 16 to 30 years^9^, leading to its progressive accumulation and persistent toxicity. Chronic low-dose exposure has been strongly associated with pulmonary disorders, including emphysema, asthma, and bronchitis. With prolonged exposure, Cd toxicity extends far beyond the respiratory system, contributing to the development of multiple malignancies such as prostate, bladder, pancreatic, kidney, and breast cancers^10^. Long-term cadmium burden has also been implicated in osteomalacia, Itai-itai disease, cardiovascular dysfunction, and neurodegenerative disorders^11^. Furthermore, cadmium readily crosses the placental barrier, thereby exerting teratogenic effects and posing significant risks to fetal development ^12^. The liver, as the primary detoxifying organ, is particularly vulnerable since Cd enters via contaminated food and water, where Cd²⁺ binds to thiols, depletes antioxidants like glutathione, which in turn increases oxidative stress that disrupts detoxification pathways and causes fibrosis and necrosis via ROS overproduction^13^. Similarly, the lungs are critically affected as Cd is present in air pollution and cigarette smoke, enabling inhalation exposure that triggers acute inflammation transitioning to emphysema and fibrosis through cytokine release (TNF-α, IL-6) and persistent ROS generation, elevating lung cancer risk^9,14^.

At the molecular level, many of these toxic effects are generated due to the strong affinity of Cd(II) for sulfur-containing biomolecules. Cd(II), as a soft Lewis acid, forms coordinate covalent bonds with thiol sulfur, a soft Lewis base from cysteine residues in glutathione and metallothionein^15^. This binding depletes antioxidants, induces oxidative stress, disrupts hepatic detoxification, and causes liver fibrosis and necrosis^16^, while inhaled cadmium triggers pulmonary inflammation via reactive oxygen species (ROS) and cytokines^17,18,19^. Cd (II) also exerts its deleterious effects partly through disruption of cellular metal homeostasis^20^. Although Cd(II) (*d¹⁰* closed-shell) and Cu(II) (*d⁹*) exhibit distinct electronic configurations, their shared affinity for soft thiolate (Cys) and imidazole (His) ligands enables Cd(II) to compete for Cu(II)-binding sites in metalloproteins such as P1B-ATPases (e.g., Cad A) and metallothioneins, often with micromolar affinities^21,22^. Analysis of Protein Data Bank structures reveals Cd(II) substitutes into Cu(II)/Zn(II) sites without major conformational disruption, supported by comparable interaction energies and side-chain preferences^23,24^. In this study, we have asked whether biological systems exploit this molecular mimicry, harnessing copper exporters to actively detoxify cadmium.

As an essential micronutrient, copper is required for a wide range of physiological processes in virtually all cell types^25^. Because the accumulation of intracellular copper can induce oxidative stress and perturb cellular function, copper homeostasis is tightly regulated^25,26^. A key player of this regulatory network is the family of Copper-transporting P(IB)-type ATPases, which are highly conserved in aerobic organisms and function to deliver copper to cupro-enzymes in the secretory pathway, transport copper across epithelial and endothelial cell membranes and efflux excess copper^27^. Interestingly, while unicellular eukaryotes and invertebrates utilise one copper-transporting P(IB)-type ATPase, a gene duplication has occurred during vertebrate evolution, resulting in *ATP7A* (*MNK*) and *ATP7B* (*WND*)^28^. Orthologs of both of these genes are present in chicken and fish species, suggesting the duplication occurred early in chordate evolution^29^. The human paralogues (*ATP7A*: NM_000052.7; *ATP7B*: NM_000053.4) share approximately 54 percent sequence identity but differ in their tissue expression profile^30^. ATP7A is expressed almost ubiquitously with the exception of the liver, whereas ATP7B is predominantly expressed in the liver as well as the brain, kidney, placenta and mammary glands^31^. Mutations in *ATP7A* are responsible for the systemic copper deficiency in Menkes disease due to copper accumulation in the gut and kidney^32^, whereas mutations in *ATP7B* cause Wilson disease with copper toxicity due to accumulation in the liver^33^. The ATP-driven copper pumps (Copper-ATPases) play a critical role in living organisms by maintaining optimum copper levels in cells and tissues. Among the key functional domains of Copper-ATPases, the metal-binding N-terminal domain could be responsible for the functional diversification of the copper ATPases during the course of evolution^29^.

The human ATP-dependent copper exporter ATP7B holds a significant position in maintaining cellular copper homeostasis. It transports intracellular copper in two ways. Usually, ATP7B is located in the *trans*-Golgi network and delivers copper into the Golgi for the biosynthesis of copper-containing enzymes (cuproenzymes). When the cytosolic copper concentration elevates, ATP7B traffics to lysosomes, facilitating copper excretion into bile along with vesicle fusion^34^. In addition, overexpression of ATP7B is relevant to resistance to several chemotherapeutic agents, such as cisplatin^35^ and ruthenium, in tumour cells^36^. To elucidate the phenomenon, some investigations have shown that cisplatin can be sequestered by the CXXC motifs of ATP7B MBDs or may undergo an ATP-dependent transport similar to Cu^+^ and become sequestrated into endo-lysosomal compartments^37^.

In our study, we found that cadmium induces trafficking of both ATP7A and ATP7B from the trans-Golgi network (TGN) in response to metal overload, with ATP7B specifically colocalising to lysosomes as an adaptive detoxification strategy to sequester and export Cd(II), while the Cd-induced trafficking of ATP7A in lung cell lines remains to be elucidated. In parallel, *C. elegans* models demonstrated that the copper-transporting P1B-ATPase cua-1 and lysosome-related organelles (LROs) play pivotal roles in Cd(II) detoxification.

## Results

### ATP7A undergoes vesicularization in presence of excess cadmium in lung adenocarcinoma cells

Cadmium (Cd) is a toxic heavy metal that accumulates in the lungs^38^. Disruption of metal homeostasis, particularly cellular copper homeostasis, is a key feature of Cd(II) induced toxicity. A549 lung adenocarcinoma cells predominantly express ATP7A^39^. We therefore investigated whether ATP7A exhibits vesicular and lysosomal localization in response to CdCl_2_ treatment in A549 cells.

A549 cells were plated on a coverslip and treated with excess CuCl_2_, followed by treatment with ammonium tetrathiomolybdate (TTM). Under basal condition, ATP7A exhibited colocalization primarily with the TGN marker (Golgin 97), along with negligible vesicular distribution (Fig.1A, upper panel). Upon treatment with 50 µM CuCl_2_ for 2 hours, ATP7A showed vesicularization and colocalization with TGN marker Golgin 97 (Fig.1A, middle panel), indicating ATP7A can perform Cu-induced anterograde trafficking. Following CuCl_2_ treatment for 2 hours, cells are subsequently treated with 50 µM TTM for 2 hours. This resulted in the retrograde trafficking of ATP7A from vesicular compartments back to the TGN, as evidenced by reduced vesicularization and an increase in colocalization with the Golgin 97 (Fig.1A, lower panel). Based on the findings, we established that ATP7A undergoes both anterograde and retrograde trafficking in response to CuCl_2_ and TTM treatment in A549 cells. To determine whether Cd(II) elicits a comparable trafficking pattern, A549 cells were treated to increasing concentrations of CdCl_2_ (20 µM, 50 µM, and 100 µM). Notably, ATP7A exhibited a dose-dependent increase in vesicular localization upon CdCl_2_ treatment, resembling the similar kind of vesicularization pattern observed under 50 µM CuCl_2_ treatment condition (Fig.1B). This suggests that ATP7A can perform the Cd(II) induced anterograde trafficking. Although the PCC values (Basal with 20µM CdCl_2_) did not show a statistically significant decrease however, a sequential decline in PCC (20 µM CdCl_2_ with 50 µM CdCl_2_) and (50 µM CdCl_2_ with 100 µM CdCl_2_) was observed (Fig.1C).

**Figure 1.**
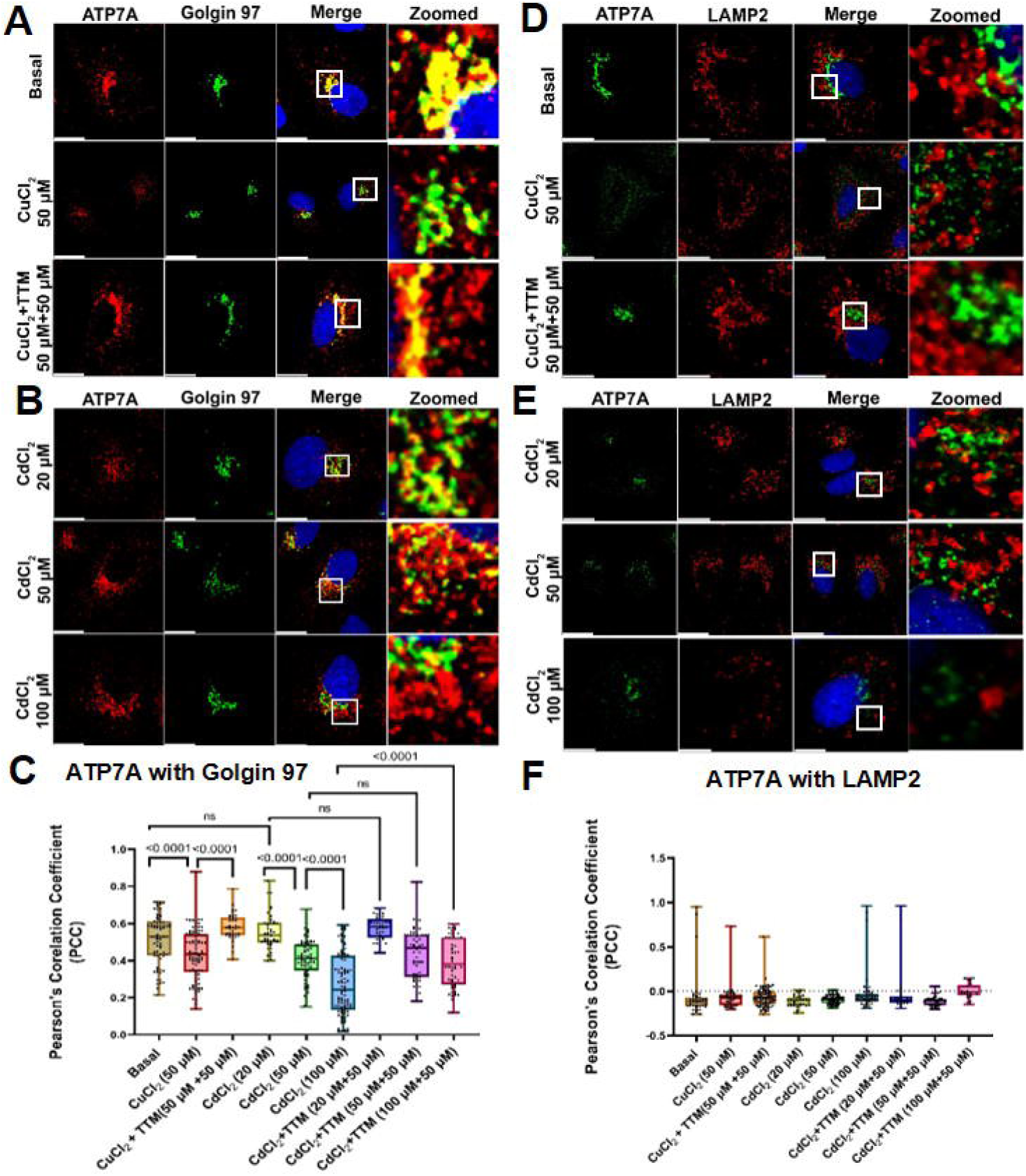
Localization of ATP7A following CuCl_2_ and CdCl_2_ treatment in A549 cells. (A) Localization of ATP7A (red) with Golgin-97 (green) under basal conditions (upper panel). ATP7A predominantly colocalizes with the Golgin-97 marker, apart from its presence in vesicular structures. Upon CuCl_2_ treatment, ATP7A undergoes vesicularization (middle panel). Following CuCl_2_ treatment and subsequent treatment with tetrathiomolybdate (TTM), ATP7A traffics back to the TGN (lower panel).(B) A549 cells were treated with 20μM, 50μM, and 100μM of CdCl_2_. ATP7A similarly undergoes vesicularization in a dose-dependent manner. (C) Pearson’s correlation coefficient (PCC) is quantified. A total of 75 cells (basal), 91 cells (50μM CuCl_2_-treated), 39 cells (20μM CuCl_2_ + 50μM TTM), 48 cells (20μM CdCl_2_ treated), 70 cells (50μM CdCl_2_ treated), 100 cells (100μM CdCl_2_ treated), 41 cells (20μM CdCl_2_ +50μM TTM treated), 60 cells (50μM CdCl_2_ +50μM TTM treated), and 54 (100μM CdCl_2_ +50μM TTM treated) were analysed. p values for each condition are indicated in the graph. (D) A549 cells were stained for ATP7A (green) and LAMP2 (red). Under basal conditions (upper panel), as well as following CuCl_2_ treatment and CuCl_2_ + TTM treatment, no significant colocalization of ATP7A with LAMP2 has been observed.(E) Similarly, treatment with 20μM, 50μM, and 100μM CdCl_2_ doesn’t not show colocalization between ATP7A and LAMP2.(F) PCC values were quantified from 38 cells (basal), 39 cells (CuCl_2_-treated), 85 cells (CuCl_2_ + TTM), 32 cells (20μM CdCl_2_ treated), 71 cells (50μM CdCl_2_ treated), 49 cells (100μM CdCl_2_ treated), 27 cells (20μM CdCl_2_ +50μM TTM treated), 45 cells (50μM CdCl_2_ +50μM TTM treated), and 65 cells (100μM CdCl_2_+50μM TTM treated). PCC analysis showed no significant colocalization between ATP7A and LAMP2 under any condition. Scale bar: 5μm

To further determine whether the observed vesicularization of ATP7A is directly induced by Cd(II), rather than being a secondary consequence of increased intracellular Cu(I) amount due to possible metabolic shift in the cell, we performed copper-chelation experiments using TTM, post-Cd(II) treatment. A549 cells were treated with increasing concentrations of CdCl_2_ (20 µM, 50 µM, and 100 µM) for 2 hours, followed by treatment with 50 µM TTM. Notably, ATP7A were retained its vesicular localization upon TTM treatment, with no retrograde trafficking to TGN (Fig.S1A and Fig.S1B). This suggests that Cd(II)-induced vesicularization of ATP7A is specific and not dependent on intracellular Cu(I) availability. Quantitative analysis using PCC further supported these observations (Fig.1C). No significant differences are observed between CdCl_2_ only (20 µM and 50 µM) and corresponding CdCl_2_+TTM-treated conditions (20 µM+50 µM and 50 µM+50 µM). However, at the higher concentration (100 µM CdCl_2_), a significant change in PCC values was observed between CdCl_2_-only and CdCl_2_+TTM-treated cells. This shift in the PCC value at 100 µM may reflect the combined effects of the increased Cd(II) exposure and TTM-induced cellular stress. Collectively, these results indicate that treatment with CdCl_2_ triggers vesicularization ATP7A in A549 cells.

To determine whether CdCl_2_ induced vesicularization of ATP7A is not cell type-specific, we extended our analysis to additional cell lines, including HEK293 (Fig.S2) and HeLa cells (Fig.S3), both of which predominantly express ATP7A as their major copper transporter. Cells were treated with excess CuCl_2_ or CdCl_2_ under conditions similar to those used for A549 cells. CuCl_2_ treatment induced vesicularization of ATP7A in both HEK293 and HeLa cells, comparable to that observed upon CuCl_2_ treatment. This suggests that Cd(II)-triggers redistribution of ATP7A is not restricted to lung-derived cells but represents a more generalized cellular response. Furthermore, copper chelation with TTM did not reverse Cd(II)-induced vesicularization of ATP7A in these cell lines, consistent with our observations in A549 cells (Fig.S2 and Fig.S3).

To further validate that ATP7A vesicularization is specifically induced by Cd(II) rather than other divalent metal ions, A549 cells were treated with, ZnCl_2_, MgCl_2_ and CaCl_2_. Notably, these treatments did not induce vesicularization of ATP7A (Fig.S4) suggesting the ATP7A vesicularization is restricted to Cu and Cd(II) elevation exclusively.

To define the identity of ATP7A-containing vesicles following CuCl_2_/ CdCl_2_ treatment, we investigated their colocalization with the lysosomal marker LAMP2 to determine whether ATP7A facilitates the lysosomal exocytosis to export cadmium. Under excess CuCl_2_ conditions, ATP7A did not colocalize with LAMP2-positive vesicles, indicating that ATP7A is unlikely to perform copper export into lysosomes in A549 cells (Fig. 1D). Consistent with this observation, ATP7A also failed to colocalize with LAMP2-positive lysosomal compartments upon CdCl_2_ treatment in a dose-dependent manner (20 µM, 50 µM, and 100 µM) (Fig. 1E). Furthermore, copper chelation with TTM did not alter ATP7A localization with respect to LAMP2-positive compartments (Fig.S1B). Quantitative analysis using PCC confirmed no colocalization between ATP7A and LAMP2 under all experimental conditions (Fig.1F). Collectively, these findings demonstrate that neither Cu(II)-nor Cd(II)-induced vesicularization of ATP7A corresponds with lysosomal exocytosis. Instead, ATP7A likely redistributes to non-lysosomal vesicular compartments, suggesting an alternative pathway for metal handling in A549 cells.

To identify the ATP7A-positive vesicles, we investigated the colocalization of ATP7A with components of the canonical TGN exit and late secretory trafficking pathways. Specifically, we performed the colocalization with markers of early endosomes (EEA1), late endosomes (RAB7), late recycling endosomes (RAB11), the core retromer complex (VPS35), and the clathrin-associated adaptor protein complex AP-3 (Fig.S5, Fig.S6, Fig.S7). Surprisingly, ATP7A did not show detectable colocalization with any of these markers under conditions of excess CuCl_2_ or CdCl_2_ treatment. This observation suggests that Cu(I) or Cd(II)-induced ATP7A vesicles do not correspond to canonical late secretory or retromer-mediated trafficking compartments.

Inductively Coupled Plasma Mass Spectrometry (ICP-MS) measurements showed dose-dependent increase of intracellular cadmium in A549 cells (Fig.S8).

### ATP7B traverses the retromer-lysosomal positive compartment but bypasses Rab7 compartments upon cadmium treatment

Since dietary ingestion of Cd-contaminated leafy vegetables is a major route of exposure^40^, and the liver is the primary organ responsible for metal detoxification and accumulation, we next investigated the effect of Cd exposure in hepatic cells. HepG2 cells, a well-established liver-derived cell line expressing the primary copper transporter ATP7B, have been used for further study.

As previously reported, a major fraction of ATP7B localizes to TGN under basal condition in HepG2 cells^42^ and consistent with this, we observed that under basal condition, ATP7B is predominantly present at the TGN (Fig.2A, top panel). Treatment with 50µM CuCl_2_ for 2 hours induces vesicularization of ATP7B, accompanied by a loss of colocalization with the TGN marker p230 (Fig. 2A, middle panel) indicating Cu-induced anterograde trafficking. Subsequent treatment with 50µM TTM following CuCl_2_ exposure restored ATP7B localization to the TGN, as evidenced by a reduction in vesicular distribution and an increase in colocalization with p230 (Fig. 2A, bottom panel), consistent with retrograde trafficking upon copper chelation.

**Figure 2.**
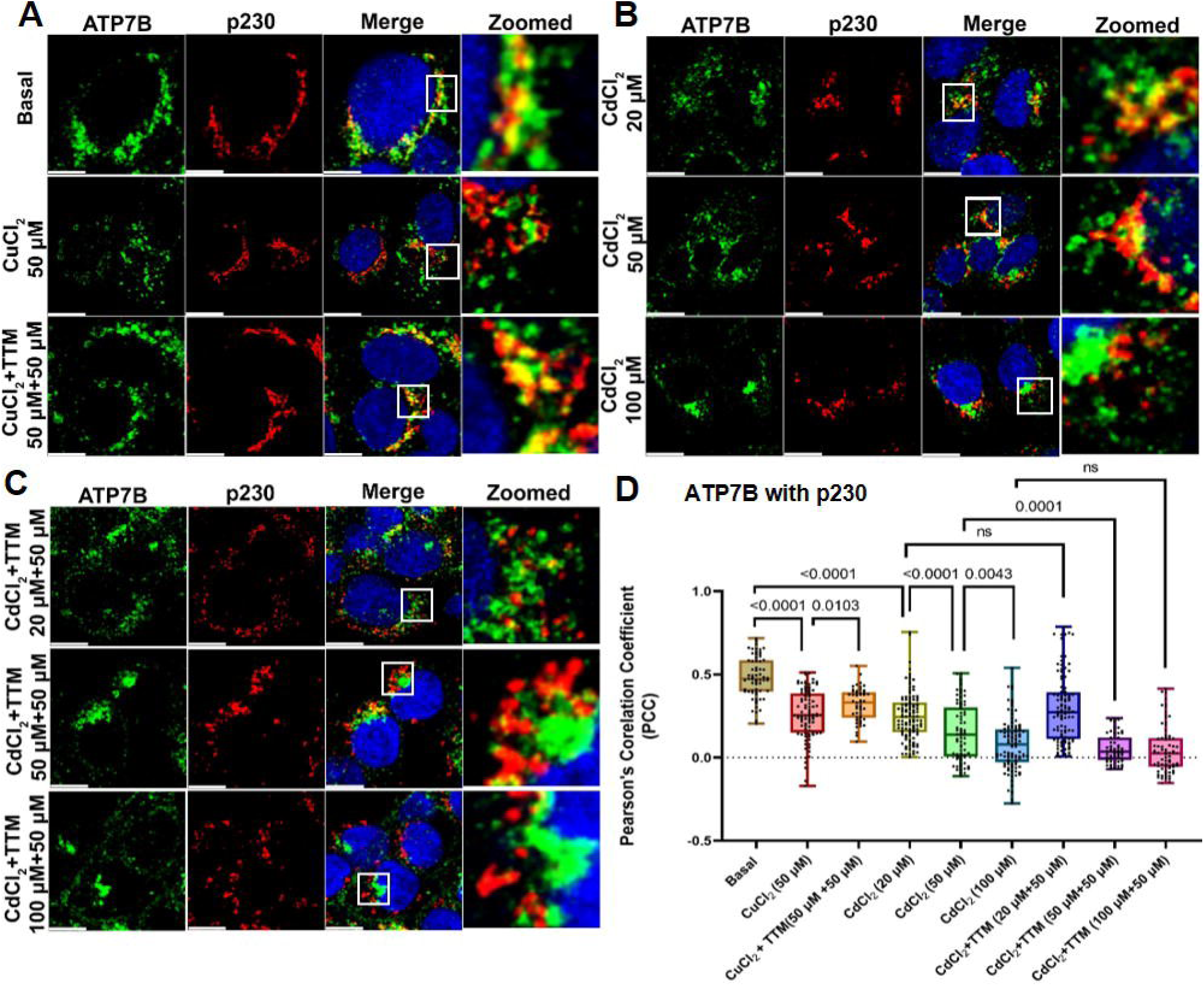
Localization of ATP7B following CuCl_2_ and CdCl_2_ treatment in HepG2 cells. (A) Localization of ATP7B (green) with p230 (red) under basal conditions (upper panel). ATP7B predominantly colocalizes with the p230 marker, apart from its presence in vesicular structures outside TGN. Upon CuCl_2_ treatment, ATP7B undergoes vesicularization (middle panel). Following CuCl_2_ treatment and subsequent treatment with TTM, ATP7B traffics back to the TGN (lower panel).(B) HepG2 cells were treated with 20μM, 50μM, and 100μM CdCl_2_. ATP7B similarly undergoes vesicularization in a dose-dependent manner. (C) HepG2 cells were treated with 20μM, 50μM, and 100μM CdCl_2_ followed by treatment with 50μM TTM. Since TTM does not chelate Cd(II), ATP7B retained a similar vesicular localization pattern in both the presence and absence of TTM treatment. (D) PCC values are quantified from 65 cells (basal), 91 cells (50μM CuCl_2_-treated),49 cells (50μM CuCl_2_ +50μM TTM treated), 113 cells (20μM CdCl_2_ treatment), 70 cells (50μM CdCl_2_ treated), 100 cells (100μM CdCl_2_ treatment), 100 cells (20μM CdCl_2_ +50μM TTM treated), 54 cells (50μM CdCl_2_ +50μM TTM treated), and 75 cells (100μM CdCl_2_ +50μM TTM treated). p values for each condition were indicated in the graph. Scale bar: 5μm

HepG2 cells treated with increasing concentrations of CdCl_2_ (20 µM, 50 µM, and 100 µM) showed a dose-dependent increase in vesicular localization of ATP7B, resembling the pattern observed upon CuCl_2_ treatment. This suggests that Cd(II) can induce the anterograde trafficking of ATP7B in HepG2 cells (Fig. 2B).

To further determine whether the observed vesicularization of ATP7B is directly induced by Cd(II), rather than being a secondary consequence of increased intracellular Cu(I) amount, we performed chelation experiments using TTM as performed in A549 cells. HepG2 cells were treated with increasing concentrations of CdCl_2_ (20 µM, 50 µM, and 100 µM) for 2 hours, followed by incubation with 50 µM TTM. Notably, ATP7B retained its vesicular localization upon TTM treatment, with no retrograde trafficking to TGN (Fig.2C).

Quantitative analysis of PCC revealed a significant decrease under 20 µM CdCl_2_ treatment compared to basal conditions. Furthermore, exposure to 50 µM and 100 µM CdCl_2_ resulted in a progressive reduction in PCC values between ATP7B and the TGN marker p230, consistent with increasing vesicular redistribution of ATP7B (Fig.2D). No significant differences are observed between CdCl_2_ only (20 µM and 100 µM) and corresponding CdCl_2_+TTM-treated conditions (20 µM+50 µM and 100 µM+50 µM). However, at the 50 µM concentration (50 µM CdCl_2_), a significant change in PCC values was observed between CdCl_2_-only and CdCl_2_+TTM-treated cells (Fig. 2D). Collectively, these results indicate that ATP7B can be vesicularized in presence of CdCl_2_ in a dose dependent manner.

It has been well established that under basal conditions ATP7B localizes to TGN-lysosome contact sites ^41^, and substantial evidence indicates that ATP7B traffics to lysosomes upon exposure to excess Cu(I) in HepG2 cells^35,43^. Consistent with these reports, we observed ATP7B localization with LAMP2-positive compartments under basal conditions, as well as following treatment with 50 µM CuCl_2_ and 50 µM CuCl_2_+ 50 µM TTM treated condition (Fig.3A), indicating copper-dependent lysosomal trafficking. Notably, treatment with increasing concentrations of CdCl_2_ resulted in a dose dependent increase in colocalization of ATP7B with the lysosomal marker LAMP2 (Fig.3B).

**Figure 3.**
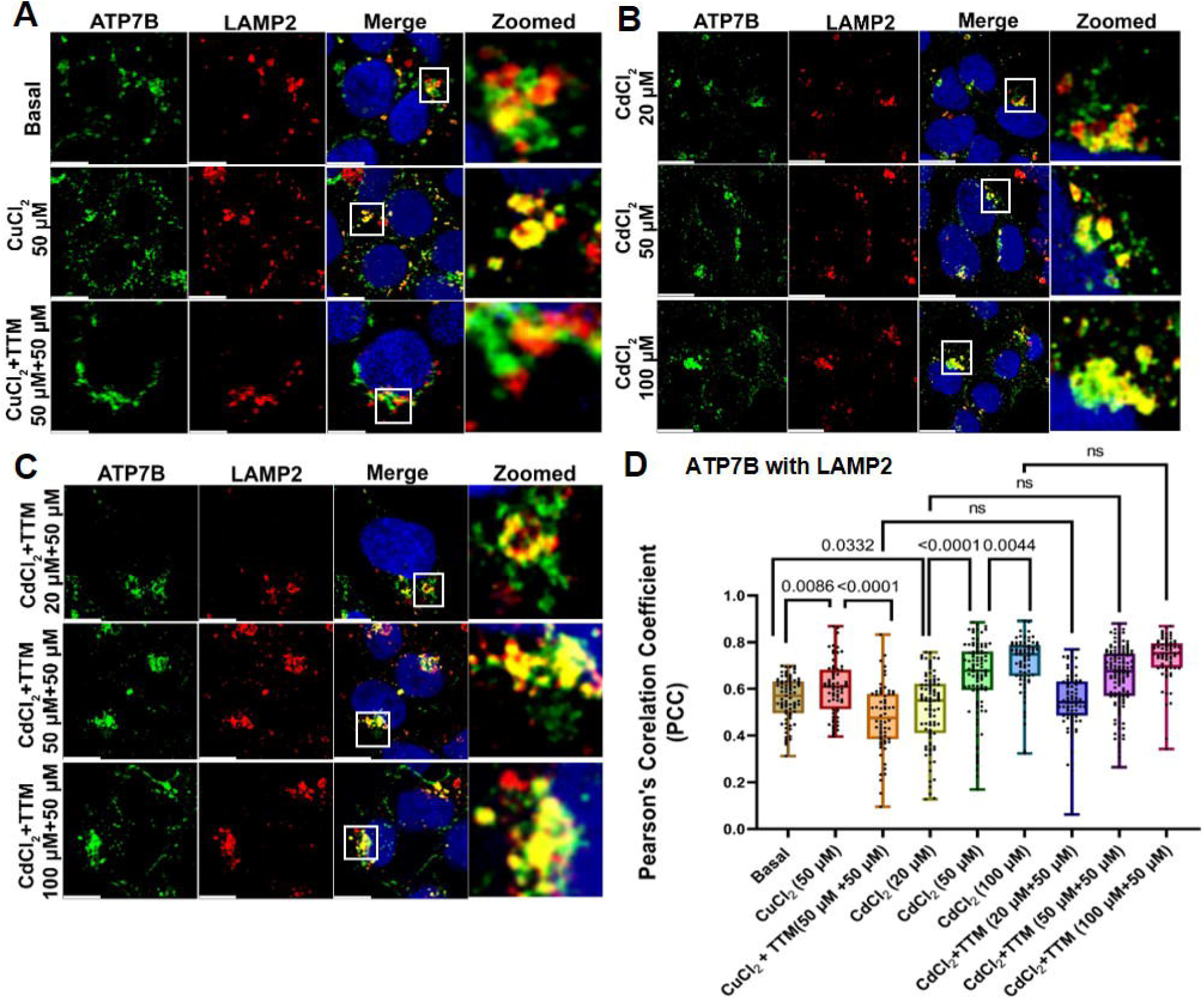
Localization of ATP7B in lysosome following CuCl_2_ and CdCl_2_ treatment in HepG2 cells. (A) Localization of ATP7B (green) with LAMP2 (red) under basal conditions (upper panel). A part of ATP7B colocalizes with the LAMP2 marker. Upon CuCl_2_ treatment, ATP7B undergoes predominant colocalization with LAMP2 (middle panel). Following CuCl_2_ treatment and subsequent treatment with TTM, ATP7B traffics back to the TGN. Hence less colocalization with LAMP2 has been observed (lower panel).(B) HepG2 cells were treated with 20μM, 50μM, and 100μM CdCl_2_. ATP7B similarly undergoes colocalization with LAMP2 positive lysosomal compartment in a dose-dependent manner. (C) HepG2 were treated with 20μM, 50μM, and 100μM CdCl_2_ followed by treatment with 50μM TTM. Since TTM does not chelate Cd(II), ATP7B retained a similar LAMP2 positive vesicular localization pattern in both the presence and absence of TTM treatment. (D) PCC values are quantified from 75 cells (basal), 66 cells (50μM CuCl_2_-treated),62 cells (50μM CuCl_2_ + 50μM TTM treated), 89 cells (20μM CdCl_2_ treated), 96 cells (50μM CdCl_2_ treated), 74 cells (100μM CdCl_2_ treated), 86 cells (20μM CdCl_2_ +50μM TTM treated), 124 cells (50μM CdCl_2_ +50μM TTM treated), and 68 cells (100μM CdCl_2_ +50μM TTM treated). p values for each condition are indicated in the graph. Scale bar: 5μm

As expected, subsequent treatment with CdCl_2_+ TTM failed to relocalize ATP7B back to the TGN, suggesting that Cd(II)-induced lysosomal localization of ATP7B is independent of intracellular copper availability and cannot be reversed by copper chelation (Fig.3C). PCC analysis further supports this observation (Fig.3D), reinforcing the conclusion that Cd(II) drives a distinct lysosome-associated trafficking of ATP7B.

To further validate that ATP7B vesicularization is specifically induced by Cd(II) rather than other divalent metal ions, HepG2 cells were treated with, ZnCl_2_, MgCl_2_ and CaCl_2_. Notably, treatment with Mg(II) and Ca(II) did not induce vesicularization of ATP7B (Fig.S9)

We next investigated whether it follows a similar endocytic route as observed during ATP7B-mediated lysosomal sequestration of excess intracellular Cu(I). Previous study from our group has shown that ther retromer complex comprising of VPS35, VPS26 and VPS29 regulates trafficking of ATP7B in hepatocytes ^42^. Upon treatment with increasing doses of CuCl_2_ (20 µM and 50 µM) for 2 hours, ATP7B was found to colocalize with VPS35. A similar pattern of colocalization was also observed under CdCl_2_ treatment (20 µM and 50 µM) for 2 hours (Fig.4A and 4B). Pearson’s correlation coefficient (PCC) analysis further indicated a comparable trend (Fig. 4C).

**Figure 4.**
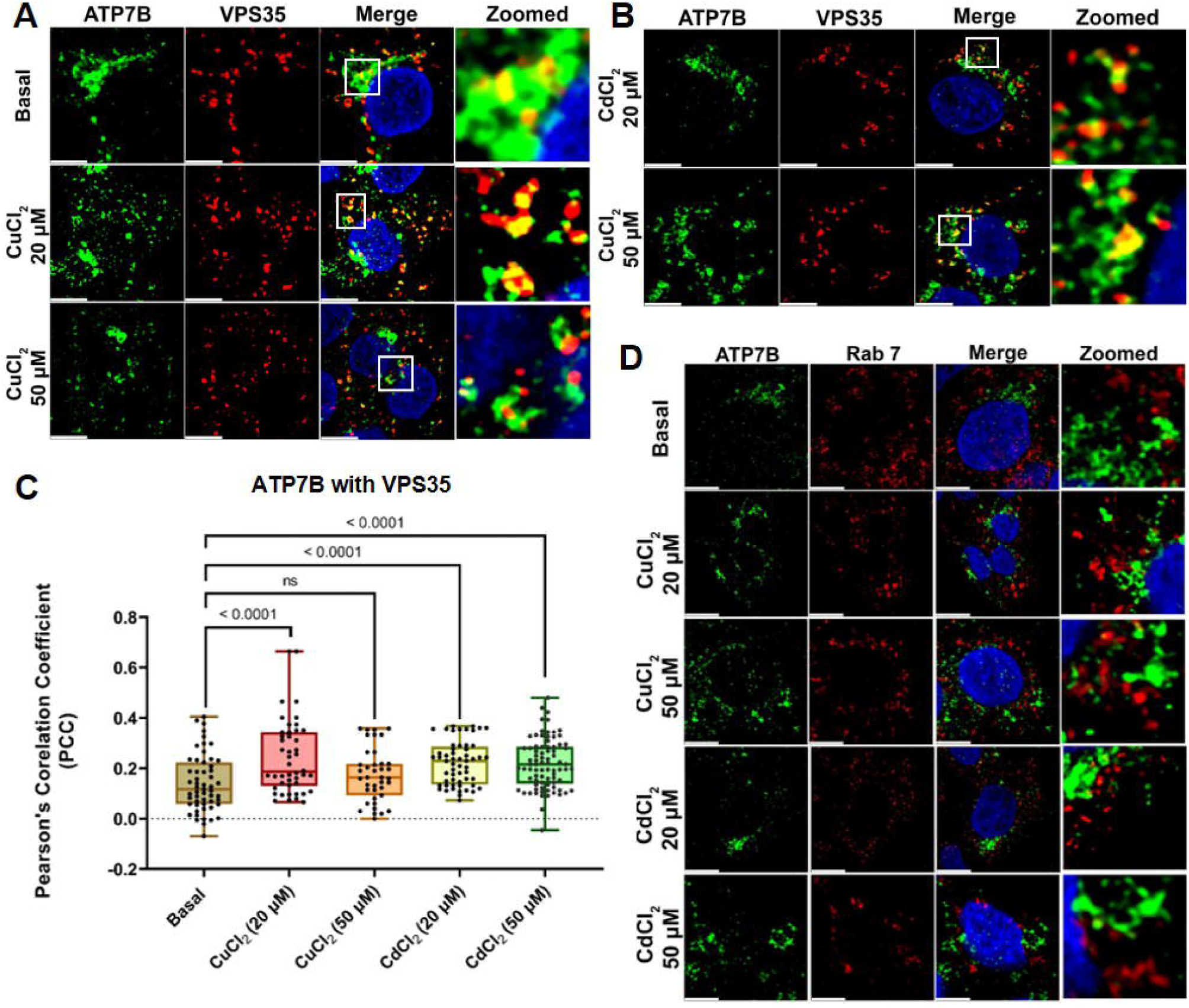
Localization of ATP7B with VPS35 and RAB 7 following CuCl_2_ and CdCl_2_ treatment in HepG2 cells. (A) Localization of ATP7B (green) with VPS35 (red) under basal conditions (upper panel). A part of ATP7B colocalizes with the VPS35 (retromer positive endosome) marker. Upon 20μM of CuCl_2_ treatment, ATP7B undergoes colocalization with VPS35 (middle panel). Upon 50μM of CuCl_2_ treatment, the colocalization of ATP7B with VPS35 has decreased (lower panel). (B) HepG2 cells were treated with 20μM and 50μM, CdCl_2_. ATP7B undergoes colocalization with VPS35 in a dose-dependent manner. (C) PCC values are quantified from 55 cells (basal), 46 cells (20μM CuCl_2_ treated),39 cells (20μM CuCl_2_ treated), 56 cells (20μM CdCl_2_ treated), 96 cells (50μM CdCl_2_ treated). (D) localization of ATP7B (green) and RAB7(red) under basal condition (first panel). No significant colocalization of ATP7B with RAB7(late endosome) has been observed. Under conditions of increased CuCl_2_ treatment i.e. 20μM (second panel) and 50μM (third panel) no significant colocalization of ATP7B and RAB7 has been observed. Similarly, treatment with 20μM (fourth panel) and 50μM CdCl_2_ (fifth panel) doesn’t not show colocalization of ATP7B with Rab7.Scale bar: 5μm

However, neither CuCl_2_ nor CdCl_2_ treatment (20 µM and 50 µM for 2 hours) resulted in significant colocalization of ATP7B with the early endosomal marker EEA1 (Fig.S10) or the late endosomal marker Rab7 (Fig.4D). This suggests that upon cadmium treatment, ATP7B bypasses the Rab7 compartment or exhibits short-time residence at late endosomal compartments before terminating at the lysosomes.

### ATP7B provides Cadmium resistance and sequesters the metal

The hepatocytic copper-transporting P-type ATPase, ATP7B, maintains copper homeostasis mainly by trafficking from the Golgi to lysosomes upon elevated copper levels ^34^. This transporter handles xenobiotics structurally similar to copper, such as platinum-based chemotherapeutics (cisplatin^35^, carboplatin^43^) and silver ions^44^. Platinum binds ATP7B’s metal-binding domains, triggering vesicular sequestration and lysosomal export, which reduces drug efficacy in cancer cells and promotes resistance. Similarly, in airway ciliated cells, ATP7B traffics apically in response to silver, aiding detoxification^37,44^.

To analyse the role of ATP7B in detoxifying Cadmium, we hypothesised that the absence of ATP7B in HepG2 cells should increase their sensitivity to Cd-induced death. Wildtype and ATP7B knockout HepG2 cells were treated with increasing concentrations of CdCl_2_ for 3 hours and 24 hours respectively. As compared to WT cells, ATP7B ^−/−^ cells exhibit higher sensitivity (Fig.5A and Fig.5B). The IC_50_ values for 3 hours and 24 hours of CdCl_2_ treatment reveals higher toxic effect of cadmium on ATP7B ^−/−^ cells (Fig.5C and Fig.5D), suggesting a potential role of Cd sequestration by ATP7B in hepatocytes.

**Figure 5.**
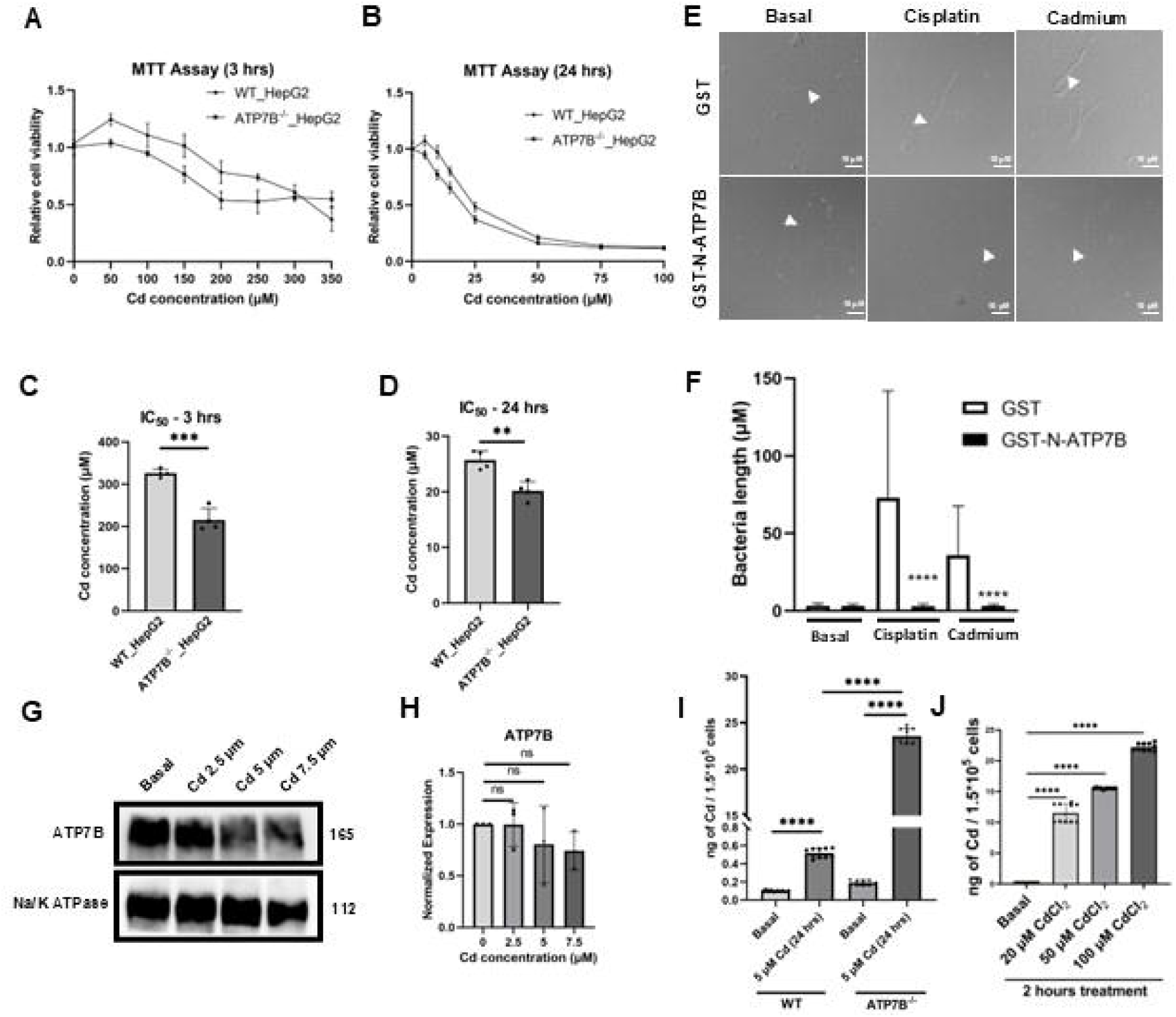
ATP7B protects HepG2 cells from cadmium-induced toxicity through metal sequestration and cellular clearance mechanisms. (A, B) MTT-based cell viability assays of WT and ATP7B^−/−^ HepG2 cells following exposure to increasing concentrations of cadmium (Cd) for 3 h (A) and 24 h (B). (C, D) Quantification of IC_50_ values derived from the viability assays shown in panels A and B after 3 h (C) and 24 h (D) Cd exposure. (E) Representative DIC confocal images of *E. coli* expressing GST alone or the GST-tagged N-terminal metal-binding domain of ATP7B (GST–N-ATP7B) under basal conditions or following treatment with cisplatin or Cd.(F) Quantification of bacteria length shown in panel E. (G) Representative immunoblot showing ATP7B protein expression in WT HepG2 cells following 24 h exposure to increasing Cd concentrations (2.5–7.5 μM). α-tubulin served as a loading control. (H) Densitometric quantification of ATP7B protein levels normalised to α-tubulin from panel G. (I) WT and *ATP7B^−/−^* HepG2 cells are treated with 5μM CdCl_2_ for 24 hours. Intracellular Cd levels were measured by ICP-MS and represented as ng Cd/1.5 × 10^5 cells. Under basal conditions, the Cd level is 0.1035 ng/1.5 × 10^5 cells and 0.1915 ng/1.5 × 10^5 cells for WT and *ATP7B−/−* HepG2 cells, respectively. Upon treatment with 5μM CdCl_2_ for 24 hours, intracellular Cd accumulation is 0.5175 ng/1.5 × 10^5 cells and 23.5545 ng/1.5 × 10^5 cells, for WT and *ATP7B^−/−^* HepG2 cells, respectively. Number of biological replicates = 2; number of technical replicates = 5 for each condition. p-values for each condition are written on the graph (J). HepG2 cells are treated with 20μM, 50μM, and 100μM CdCl_2_ for 2 hours. Intracellular Cd levels are measured and represented as ng Cd/1.5 × 10^5 cells. Under basal conditions, the Cd level is 0.207 ng/1.5 × 10^5 cells. Upon treatment with 20μM CdCl_2_, intracellular Cd accumulation increased to 11.534 ng/1.5 × 10^5 cells, while treatment with 50μM and 100μM CdCl_2_ resulted in Cd accumulation levels of 15.527 ng/1.5 × 10^5 cells and 22.242 ng/1.5 × 10^5 cells, respectively. Number of biological replicates = 2; number of technical replicates = 5 for each condition. p-values for each condition are written on the graph. In all the experiments, n≥3, and in E, F, N=100 and each data point represents one biological replicate, except mentioned otherwise. Data are presented as mean ± SD. Statistical significance was determined using an unpaired Student’s t-test with appropriate multiple-comparison analysis. **p < 0.01, ***p < 0.001, ****p < 0.0001.

Based on this experimental observation, we wanted to test whether the amino terminal harbouring six copper-binding CXXC motifs of ATP7B^45^ is able to sequester Cd(II). Based on the previous work from our group, we overexpressed GST-N-ATP7B in *E. coli* and treated the cells either with Cisplatin or Cadmium^36^. As cisplatin treatment induces the formation of bacterial filaments as a stress response^35^, we have used cisplatin as our positive control and compared Cd-induced filamentation in the control *E. coli* cells and cells overexpressing the N-ATP7B. The average length of the N-ATP7B-overexpressed bacteria was significantly lower than that of the groups expressing the GST-tag under cadmium treatments. (Fig.5E, 5F).

To further determine whether Cd exposure alters ATP7B protein pool, WT HepG2 cells were exposed to increasing concentrations of CdCl₂ (2.5 - 7.5 μM) for 24 hours, followed by western blot analysis of ATP7B expression. ATP7B protein levels remained largely unchanged across all treatment conditions, with no statistically significant differences compared to basal controls (Fig. 5G, 5H). These findings suggest that Cd detoxification is unlikely to involve translational upregulation of ATP7B under Cd stress conditions; instead may primarily depend on redistribution or trafficking of the pre-existing ATP7B pool.

To consolidate our findings, we tested whether ATP7B^−/−^ HepG2 cells accumulate more Cd compared to WT cells. For that, we exposed HepG2 cells with 5 μM CdCl2 for 24 h. Cd-treated WT cells showed Cd levels comparable to untreated controls. In contrast, ATP7B cells accumulated higher intracellular Cd(II), suggesting that ATP7B plays a critical role in limiting Cd accumulation (Fig.5I). Acute cadmium treatment also leads to its increasing accumulation in HepG2 cells in a dose-dependent manner (Fig.5J).

In summary, our findings establish ATP7B as a critical mediator of cadmium (Cd) detoxification in hepatocytes, extending its canonical role in copper homeostasis to structurally analogous heavy metals. ATP7B knockout in HepG2 cells markedly heightened Cd sensitivity, evidenced by increased cytotoxicity at both early (3 hrs) and prolonged (24 hrs) exposures, underscoring its protective function via endolysosomal trafficking. Complementary bacterial filamentation assay confirmed that the N-terminal metal-binding domain of ATP7B sequesters Cd, mirroring its established action on platinum chemotherapeutics, while non-significant changes of ATP7B protein upon Cd exposure support a trafficking-dominant mechanism over *de novo* synthesis. These results highlight ATP7B’s broader role in detoxifying xenobiotics.

### Cells mitigate cadmium toxicity by elevating lysosomal abundance and function

Lysosomes play a central role in cellular metal detoxification by serving as dynamic compartments for the sequestration and eventual removal of toxic metal ions^46^. They selectively concentrate metals from the cytosol, promote their precipitation into less reactive, insoluble forms within the acidic lumen and export the metal via lysosomal exocytosis^47^. Upon elevated cellular copper conditions, copper ATPase pump ATP7B traffics from the TGN to the lysosomal membrane. Being a cation pump, the acidic environment of the lumen facilitates the efficient functionality of ATP7B ^30^. Excess copper is then pumped into the lysosomal lumen by the lysosome-localised ATP7B. Apart from this canonical pathway, we hypothesised from the encouraging colocalization of ATP7B and LAMP2 data upon Cd treatment that the lysosome is also involved in the ATP7B-mediated Cd detoxification pathway.

To test the mitigation strategies adapted by the lysosomes upon Cd exposure, we used LysoTracker Red DND-99, which is a hydrophobic weak base and a viable fluorescent marker that selectively accumulates in acidic vesicular compartments, predominantly in lysosomes^48^. Using the lysosomotropic dye lysotracker red coupled with flow cytometric analysis, we have observed that, upon 3 hours (acute) of Cd exposure, the acidic compartment has significantly increased (Fig.6A, Fig.6B). This is possibly because of early stress-induced endocytic uptake.

**Figure 6.**
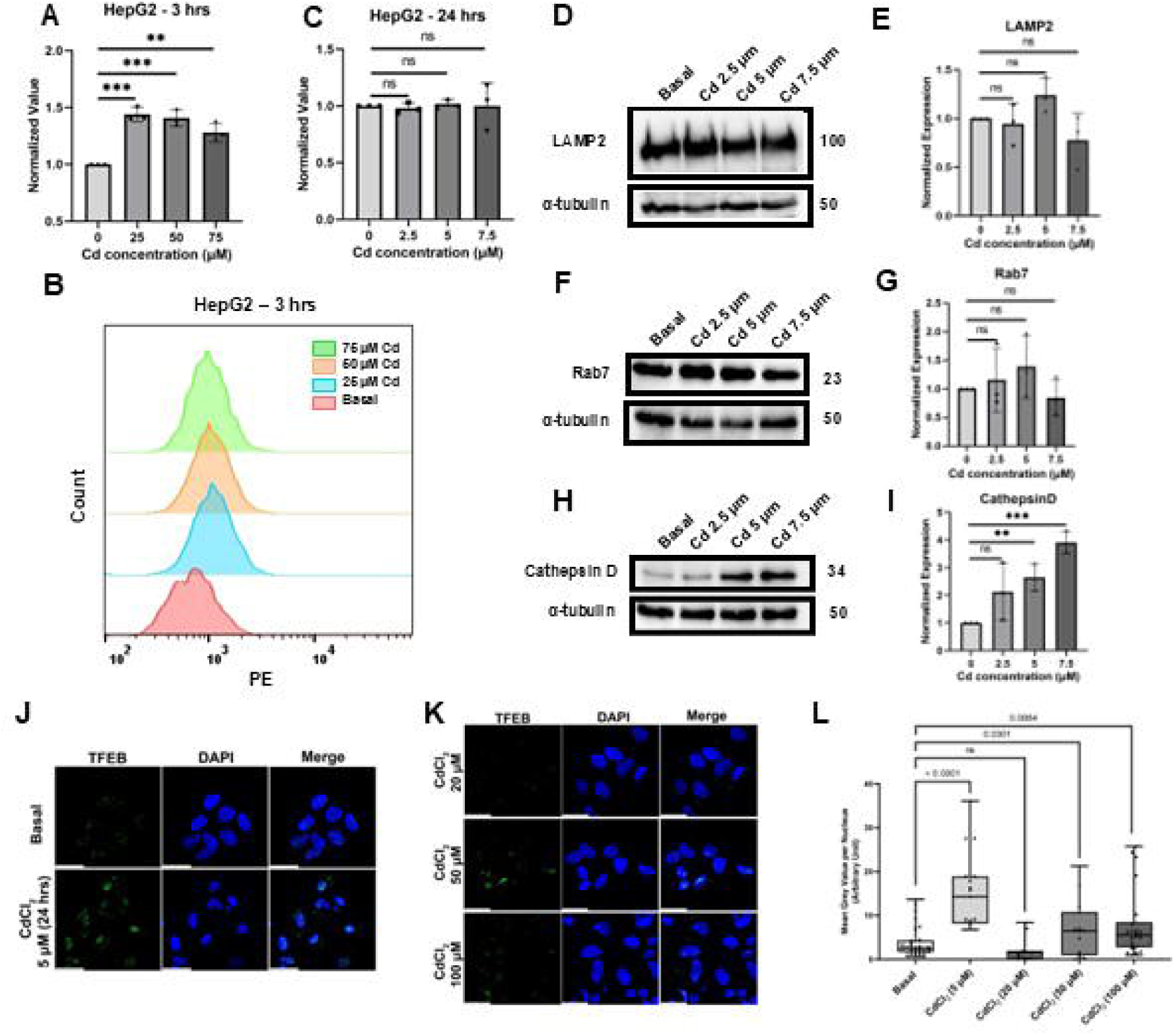
Cadmium triggers lysosomal activation and expansion of acidic vesicular compartments in HepG2 cells-. (A) Flow cytometric quantification of LysoTracker Red DND-99 fluorescence in HepG2 cells following 3 h exposure to increasing concentrations of cadmium (Cd). (B) Representative flow cytometry histograms corresponding to panel A. (C) Quantification of LysoTracker Red fluorescence after 24 h Cd exposure. (D) Representative immunoblots showing protein expression of the lysosomal membrane protein LAMP2 and late endosomal marker Rab7 following 24 h Cd exposure. α-tubulin served as a loading control. (E, F) Densitometric quantification of LAMP2 (E) and Rab7 (F) protein expression normalised to α-tubulin from panel D. (G) Representative immunoblot showing Cathepsin D expression following 24 h Cd exposure. α-tubulin served as a loading control. (H) Densitometric quantification of Cathepsin D protein levels normalised to α-tubulin from panel G. (I, J) Localisation of TFEB (green) with DAPI-stained nucleus (blue). HepG2 cells are treated with 5μM of CdCl_2_ for 24 hours. Increased localisation of TFEB in the nucleus has been observed. (K) HepG2 cells are treated with 20μM,50μM and 100μM CdCl_2_ for 2 hours. Increased localisation of TFEB in the nucleus for 50μM has been observed. (L) The nuclear signal intensity of TFEB was quantified for each condition. P values for each condition are written on the graph. Scale bar: 5μm. Each data point represents one biological replicate, except as mentioned otherwise. In every experiment, n>3. Data are presented as mean ± SD. Statistical significance was determined using an unpaired two-tailed Student’s *t*-test with appropriate multiple-comparison analysis. **p < 0.01, ***p < 0.001, ****p < 0.0001.

To further investigate lysosomal adaptation following Cd exposure, we examined the expression of key lysosomal protein markers involved in lysosomal integrity, maturation, and degradative activity. Western blot analysis revealed that the lysosomal membrane glycoprotein LAMP2 and the late endosomal/lysosomal trafficking regulator Rab7 did not exhibit significant changes following 24 hrs of Cd exposure (Fig. 6D–G), suggesting that prolonged Cd exposure does not substantially alter lysosomal abundance or late endosome–lysosome maturation at the translation level. In contrast, Cathepsin D expression was significantly elevated in a dose-dependent manner after 24 hrs of Cd exposure (Fig. 6H, I). As Cathepsin D is a major lysosomal hydrolase, its upregulation indicates enhanced lysosomal activity in response to Cd-induced cellular stress and proteotoxic burden.

We have also tested the intracellular localization of major lysosomal genes transcription factor, TFEB. We observed that TFEB preferably localizes at the nucleus upon chronic Cd treatment that is indicative of its transcriptional activity. Interestingly short-term/acute Cd treatments (20µM, 50µM and 100µM; 2 hours) we did record some TFEB nuclear localization but at a much lower level (Fig.6J. 6K and 6L).

Collectively, these findings suggest a time-dependent lysosomal response to Cadmium exposure. Acute Cd treatment induced a rapid increase in acidic vesicular compartments, as indicated by enhanced LysoTracker staining, whereas prolonged exposure activated lysosomal stress-response pathways characterised by increased Cathepsin D expression and TFEB nuclear translocation. Despite minimal changes in LAMP2 and Rab7 abundance, these results support functional lysosomal adaptation rather than a simple expansion of the lysosomal population.

### *C elegans* copper ATPase cua1 and lysosomes synergise to mitigate cadmium stress

Following the assessment of the role of copper transporter ATP7B in the cellular model, we extended our studies to an *in vivo* metazoan system, *Caenorhabditis elegans*. In *C. elegans*, the copper-exporting ATPases are represented by a single ortholog, *cua-1*, which shares sequence and functional homology with both mammalian transporters, ATP7A and ATP7B and is commonly used as an ATP7B-like model for copper-detoxification ^49^.

We used wild-type (N2 strain) and *cua1* heterozygous knockout *C. elegans* strains to compare their susceptibility towards cadmium by performing an adult survivability assay. While the wild type worms could resist up to 50µM, 5 days of Cd exposure, the number of *cua-1^−/+^* worms was significantly reduced by more than 50% upon only 10 µM Cd treatment (Fig.7A). This data supports our initial hypothesis that the copper transporting P-type ATPase has a conserved non-canonical function for detoxifying heavy metals like Cadmium, because of the absence of this particular protein caused a significant decrease in survivability in *C. elegans*. We have also checked the transcript level of *cua1*, which did not alter upon Cd exposure (Fig.7B). We measured the intracellular Cd accumulation in both N2 and *cua1^+/−^* worms following CdCl_2_ (50μM) treatment for 5 days. (Fig.7C). Cadmium levels were found to be significantly higher in *cua1^+/−^* worms as compared to the N2 strain, which further strengthens the role of cua-1 as a Cd exporter.

**Figure 7.**
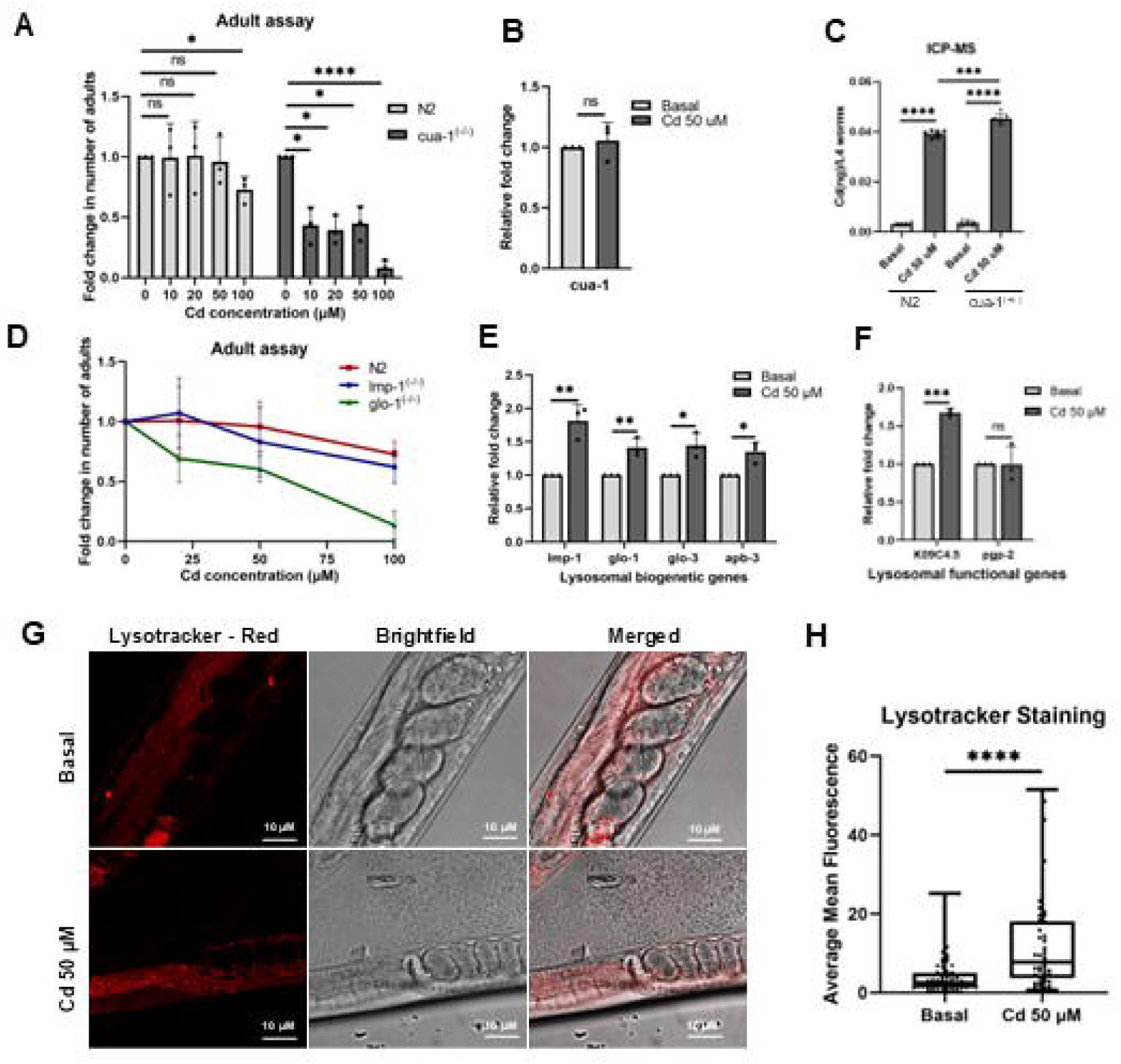
Cadmium exposure induces lysosome-related organelle responses, which promote Cd tolerance in *Caenorhabditis elegans*. (A) Adult survival assay comparing wild-type (N2) and *cua-1* heterozygous knockout worms following 5 days of exposure to increasing concentrations of cadmium (Cd). (B) Relative mRNA expression of *cua-1* in wild-type worms following exposure to 50 μM Cd. (C) ICP-MS quantification of intracellular Cd accumulation in wild-type (N2) and cua-1 heterozygous knockout worms following Cd exposure. Number of biological replicates = 2; number of technical replicates = 3 for each condition (D) Adult assay showing survival of wild-type, *glo-1^−/−^* mutant, and *lmp-1^−/−^* mutant worms following exposure to increasing Cd concentrations. Data is represented as mean ± SEM. (E) Relative expression of lysosomal and LRO biogenetic genes following 5 days of 50 μM Cd exposure. (F) Relative expression of lysosomal functional genes following Cd exposure. (G) Representative confocal microscopy images of LysoTracker Red staining in the intestinal region of *C. elegans* under basal conditions or following 50 μM Cd exposure. (H) Quantification of LysoTracker Red fluorescence intensity shown in panel G, N=30. In all experiment, n≥3, unless otherwise stated. Data are presented as mean ± SD, unless otherwise mentioned. Statistical significance was determined using an unpaired two-tailed Student’s *t*-test with appropriate multiple-comparison analysis. *p < 0.05, **p < 0.01, ***p < 0.001, ****p < 0.0001; ns, not significant.

Lysosome-related organelles (LROs), known as gut granules in *C. elegans*, share several features with canonical lysosomes, including an endosomal lineage, an acidic milieu and hydrolases, as well as lysosome-specific membrane proteins^50^. LROs extend lysosomal functions by storing metals like zinc^51^. It is reported that CUA-1 primarily localises to the basolateral membrane of the intestine under basal and copper-deficient conditions but redistributes to LROs in response to elevated copper levels in the diet ^52^.

We extended our studies to determine whether lysosomal populations play a role in mitigating Cd toxicity. To test this hypothesis, we have used *lmp-1* and *glo-1* knock-out strains, where functional lysosomes and lysosome-related organelles are absent. We found that an increased concentration of Cd increased the susceptibility of these mutants, compared to that of the wild-type *C. elegans*. (Fig.7D).

We have also checked the transcriptional regulation of lmp-1 and LRO biogenetic genes. glo-1, glo-3 and apb-3 were significantly upregulated upon 50μM, 5 day-Cd exposure (Fig.7E). The proton pump K09C4.5 localised in the gut granules also increased (Fig.7F) upon Cd exposure. This led us to test the acidic lysosomal population of *C. elegans*. As hypothesised, we indeed found a significant increase in the mean fluorescence intensity of LysotrackerRed positive lysosomal compartments upon Cd exposure in N2 worms (Fig.7G,7H).

These results together suggest that Cd exposure engages a transcriptional response involving specific LRO-associated biogenesis and transport genes, an increased acidic lysosomal population, and tolerance against Cd toxicity. Altogether, (i) the upregulated expression of genes supporting lysosomal biogenesis (ii) increased proton pump in LRO and (iii) elevated acidic lysosome population indicate a coordinated stress response against Cd exposure.

## Discussion

The canonical role of copper-transporting ATPases in maintaining Cu-homeostasis is well established^30^, but their potential role in the detoxification of non-essential toxic metals remains less explored. Although ATP7B has been implicated in the efflux of platinum-based compounds^35^, which in turn leads to chemoresistance in cancer, its role in Cd detoxification has not been studied. Here, our findings identify ATP7B as a hitherto unknown player of the cellular mitigation strategies to Cd stress.

In our studies, we show that both ATP7A and ATP7B undergo trafficking from the trans-Golgi network (TGN) upon Cd exposure, suggesting that Cd, despite being a non-essential toxic metal, engages elements of the canonical copper-responsive trafficking machinery. Cd(II), being a soft Lewis acid, has its strong affinity for cysteine-rich metal-binding domains^53^. Due to this HSAB preference, it is plausible that Cd directly interacts with the N-terminal metal-binding motifs of ATP7B, which have evolutionarily conserved CXXC motifs^37^. Consistent with this, our bacterial sequestration assays indicate that the N-terminal region of ATP7B can bind Cd and mitigate Cd-induced toxicity to bacterial cells, supporting a direct role for ATP7B in cadmium sequestration.

Our studies in mammalian cells further showed that ATP7B deficiency results in increased susceptibility to Cd exposure, which validates the protective role of ATP7B in Cd detoxification. Mechanistically, our trafficking and colocalization analyses indicate that ATP7B is redistributed to LAMP2-positive compartments upon cadmium exposure, which suggests involvement of the endo-lysosomal system. This is consistent with the established role of ATP7B in copper detoxification^30^, where it relocalizes from the TGN to vesicular compartments to facilitate metal sequestration and export^34^.

Further, acute Cd exposure of 3 hours induced a rapid increase in LysoTracker-positive acidic vesicular compartments, indicating an early lysosomal stress response prior to substantial TFEB activation. In contrast, prolonged Cd exposure of 24 hours promoted stronger TFEB nuclear translocation without significant changes in LysoTracker intensity, LAMP2, or Rab7 protein levels, suggesting transcriptional lysosomal adaptation rather than a simple increase in lysosomal abundance or acidification.

We further extended our study of Cd detoxification in the ATP7B-lysosome axis in an *in vivo* metazoan model using *Caenorhabditis elegans*. In *C. elegans* also, the Cu transporting ATPase plays an important role in Cd detoxification, supported by our adult assay data, where the absence of the ATP7 homolog, *cua-1*, makes them more susceptible towards Cd. We have also observed that cadmium exposure induced lysosomal activation and upregulation of lysosome-associated genes. Notably, we also observed induction of genes associated with lysosome-related organelles (LROs), which are the specialised compartments known to participate in metal storage and detoxification in *C. elegans*^51,54^. These findings suggest a previously unappreciated role for LRO in Cd detoxification and extend the functional landscape of lysosome-related compartments in organismal adaptation to metal stress.

Together, our findings support a model in which ATP7B contributes to Cd detoxification through a coordinated trafficking and lysosomal response (Fig.8). In this model, Cd binds to the N-terminal metal-binding domains of ATP7B, triggering its redistribution from the TGN to lysosomal compartments, where Cd is sequestered and subsequently exported. This process is coupled to nuclear localisation of TFEB, which drives transcriptional enhancement of lysosomal machinery, thereby promoting cellular adaptation to mitigate Cd stress.

**Figure 8.**
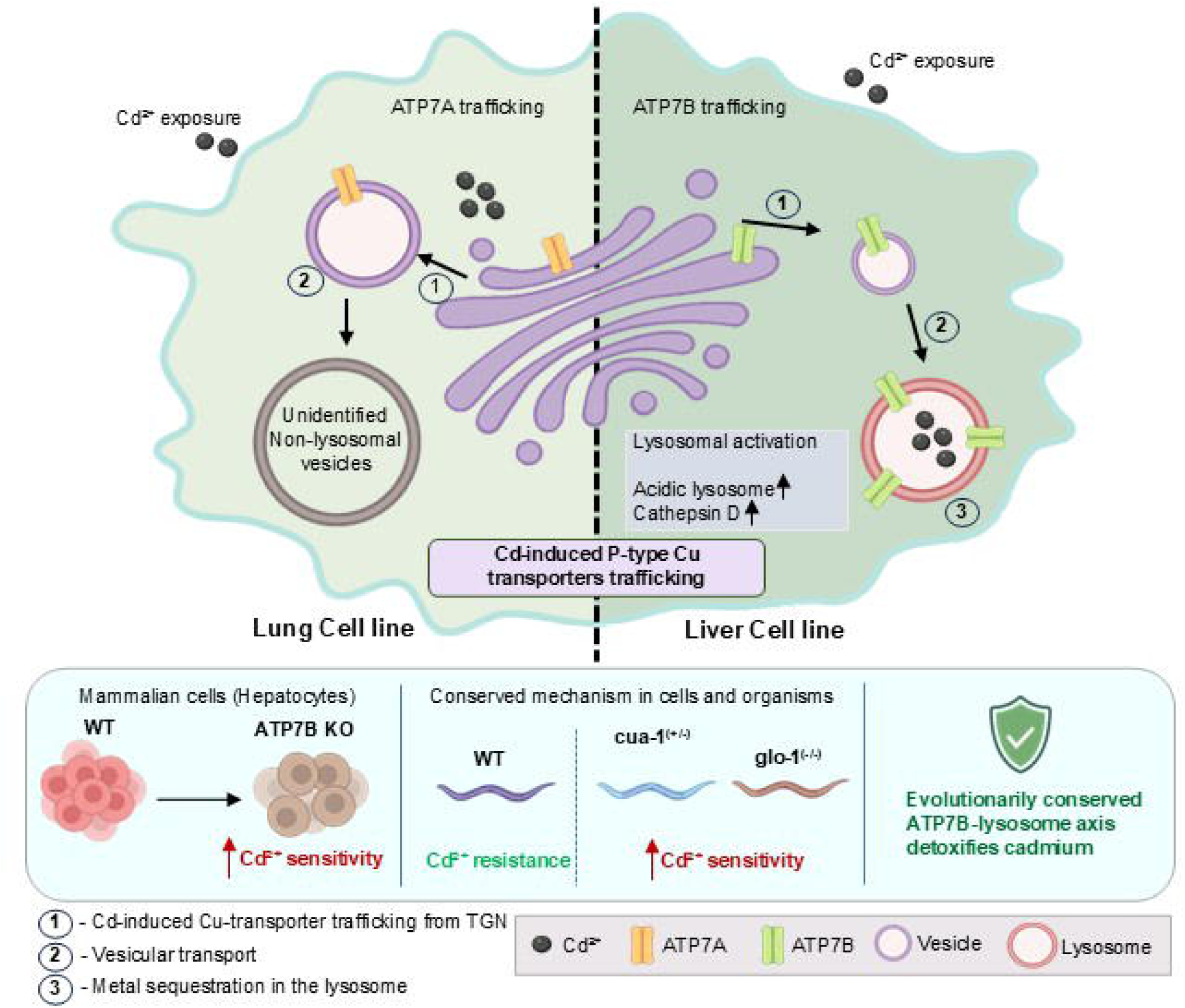
A consolidated model depicting Cadmium detoxification in lung and liver cells and *Caenorhabditis elegans*.

While these findings establish a functional link between ATP7B-lysosome machinery and Cd detoxification, several questions remain to be studied further. In particular, whether Cd directly interacts with the conserved GMXCXXC metal-binding motifs in the N-terminal domain of ATP7B, as well as the detailed trafficking itinerary and export mechanisms involved, will require further investigation. Nonetheless, our study reveals a non-canonical role for ATP7B in toxic metal handling and highlights lysosomal remodelling as a central adaptive strategy in the cellular response to cadmium toxicity.

## Experimental procedures

### Cell lines and cell culture

HepG2 cells were cultured and maintained in complete medium containing low-glucose Minimum Essential Medium (MEM) (Himedia #AL047S) supplemented with 10% Fetal Bovine Serum (Gibco #10270-106) and 1X Penicillin–Streptomycin (Gibco #15140-122).A549 cells were cultured and maintained in Dulbecco’s Modified Eagle Medium (DMEM) (Invitrogen #11995073) supplemented with 10% Fetal Bovine Serum and 1XPenicillin–Streptomycin.

For cadmium and copper treatment experiments, 10 mM stock solutions of CdCl_2_ and CuCl_2_ were prepared and used under the respective experimental conditions. To generate copper-deprived conditions, cells were treated with ammonium tetrathiomolybdate (TTM). A 10 mM TTM stock solution was prepared by dissolving the appropriate quantity of the compound in DMSO.

### Immunofluorescence and microscopy

For fixed-cell imaging, cells were fixed with 2% paraformaldehyde (PFA) in PBS for 20 min, followed by quenching of residual PFA using 50 mM NH_4_Cl solution for 20 min. Blocking was performed using 3% bovine serum albumin (BSA) prepared in PBSS (0.075% saponin in PBS). Cells were incubated with primary antibodies for 1.5 h at room temperature, followed by incubation with secondary antibodies for 30 min. Antibodies were diluted in 1% BSA prepared in PBSS.

After incubation, cells were washed three times with PBSS followed by three washes with PBS. Coverslips were then mounted onto glass slides using Fluoroshield™ with DAPI mounting medium (Sigma #F6057). Imaging was performed using a Leica Microsystems SP8 confocal microscope equipped with a 63X oil-immersion objective lens (NA 1.4). Acquired images were further deconvoluted using Leica Lightning software. The list of antibodies used in this study is provided in Table S1

### MTT assay and formazan quantifications

3-(4,5-dimethylthiazol-2-yl)-2,5-diphenyl-2H-tetrazolium bromide (MTT) is used to evaluate the toxicity of Cadmium for all the experiments. Viability was calculated on the activity of NAD(P)H-dependent cellular oxidoreductases, which reduce the MTT dye into an insoluble formazan, which is purple and can be subsequently dissolved in dimethyl sulfoxide(DMSO) for colorimetric measurement. The cells were seeded into flat-bottomed 96-well culture plates (5000 cells/200 μl/well for the WT_HepG2 and 6000 cells/200 μl/well for the ATP7B^(−/−)^_HepG2 cell line) and incubated in 37 °C, 5 % CO_2_ incubator for 72 h to allow the adherent cells to attach to the wells and let them grow up to 60–70 % confluency. Treatment was given according to the experimental requirement and incubated for 24 h. After gently pipetting off media, MTT (dissolved in PBS) was added to the MEM media (0.1 mg/ml), and the cells were incubated at 37 °C, 5 % CO_2_ for 4 h. The media was removed, and dark blue Formazan crystals formed by the cells were dissolved using 200 μl of DMSO. The absorbance was read at 570 nm wavelength on a multi-well plate reader. Data quantifications were done keeping relative viability with respect to the controls.

### Bacterial sequestration assay

BL21 (DE3) *Escherischia coli* cells were inoculated from an overnight culture in a 1:100 (v/v) ratio into 5 mL of LB broth (HiMedia) in glass test tubes. They were grown in a BOD incubator for 30 min to reach the log phase. All of the treatments were performed for 6 hrs. Smears of Cd and CDDP-treated *E. coli* cells were made on clean slides and heat-fixed. Coverslips were fixed on glass slides using SIGMA Fluoroshield™ with DAPI mountant. (Sigma #F6057). Images were taken in Leica SP8 microscope using a trans photomultiplier tube (PMT) sensor with a 63× oil immersion objective.

### Immunoblotting

After the respective CdCl_2_ treatment, cells were pelleted down. Pellet was dissolved in 100 μL of RIPA lysis buffer (10 mM Tris-Cl, pH 8.0, 1.0 % Triton X-100, 1 mM EDTA, 0.5 mM EGTA, 0.1 % sodium deoxycholate, 0.1 % SDS, 140 mM NaCl, 1X protease inhibitor cocktail, 1 mM phenylmethylsulfonyl fluoride) and kept for 45 min on ice. The solution is then sonicated with a probe sonicator (6 pulses, 5sec pulse on followed by 25 s off,100 mA). Bradford reagent (Cat #B6916, Merck) was used for protein quantitation, and 20 μg protein was loaded in each well. Protein sample was prepared by adding 4X NuPAGE loading buffer (Invitrogen#NP0007) to a final concentration of 1X and ran on 10 % SDS PAGE (8 % for ATP7B). This was further followed by semi-dry transfer of proteins onto a nitrocellulose membrane (Millipore #IPVH00010). After transfer, the membrane was blocked with 3 % BSA in TBST (TBS with 0.1 % Tween 20) for 3 h at RT with mild shaking. Primary antibody incubation was done overnight at 4 °C and then washed with 1X TBST (0.01 % Tween-20). Secondary antibody incubation was done for 2 h at RT, followed by TBST (3 times) and TBS (3 times) wash, and signal was developed by Clarity Max Western ECL Substrate (BioRad Cat #1705062) in ChemiDoc (BioRad).

### Sample preparation for ICP-MS

HepG2 and A549 cells seeded on 60 mm dishes were subjected to the respective treatment conditions at 70 % confluency. Cells were then scraped with a cell scraper after being rapidly cleaned six times with 1X DPBS (Gibco #14200075). Cells were collected by centrifuging for 5 min at 420×*g*. For counting of cells, the cell pellet was dissolved in 1 ml of 1X DPBS and counted using Countess 3 Automated Cell Counters (Invitrogen). 2.5 million cells were taken and digested overnight at 95 °C in 100 μl of ICP-OES grade 65 % HNO_3_ acid (Merck #1.00441.1000). 5 mL of 1X DPBS was added to each sample after digestion, and the samples were filtered through a 0.22-μm filter. Cadmium concentration was determined using an XSERIES 2 ICP-MS (Thermo Scientific).

For *C. elegans* cadmium measurement, a synchronised population of L4-stage worms was grown on OP50-seeded respective Cd treated 35 mm NGM plates for 5 days. 30, L4 worms were then collected in 40 μL MilliQ ultrapure water. 100 μL of Nitric acid was added to the worms, and the samples were digested in a heating block set at 95 °C overnight. The digested samples were diluted with 5 mL of MilliQ ultrapure water. Sample solutions were then thoroughly mixed and analysed. Values were normalized per L4 worm.

### Flow cytometry Assay

At 70% confluency, HepG2 cells, seeded on 24 well plates were treated with respective treatments. After treatment, cells were washed in 1X DPBS. Lysosome was stained with 100 nM LysoTracker red DND 99 (Thermo #L7528) and cells were incubated for 30 min. Cells were washed with 1X DPBS and kept in 80 μL Trypsin for 5 min. After that, cells were suspended in 400 μL of FACS buffer (1XDPBS, 2% FBS, 25mM HEPES, 2mM EDTA) and was analysed by flow cytometry BD LSRFortessa (BD Biosciences). 10,000 cells were analysed for each condition. FCS Express software (version7.14.0020) was used for data analysis. Mean intensity value were used for quantification.

### *C. elegans* strains and culture conditions

Wild-type animals used in this study were *Caenorhabditis elegans* Bristol strain N2. The *cua-1*^(+/−)^ (tm12763), *glo-1*^(−/−)^(tm3240), and *lmp-1*^(−/−)^(5826) mutant strains were obtained from the National Bioresource Project (Mitani lab, Tokyo, Japan). Worms were maintained using standard methods (Brenner) with minor modifications on nematode growth media (NGM) agar plates seeded with *Escherichia coli* OP50 as a food source. Worms were cultured at 20°C, and synchronised populations were prepared as required for experiments.

### *C. elegans* adult assay

To evaluate Cd toxicity in the WT and mutant strains, the animals were grown in NGM plates supplemented with CdCl_2_ at different concentrations ranging from 10 to 100 μM. 5 synchronised L4 hermaphrodites from WT and mutant strains were picked up and transferred singly onto a plate with or without cadmium and were grown for 5 days at 20 °C. The total number of adults was counted after 5 days. For each treatment condition, the number of adult worms was normalised to the number of adults on the basal NGM plate of that particular strain.

### quantitative Real-time PCR

Worms were collected from 35mm NGM plates after 5 days long respective treatment using 1 ml of M9 buffer and washed 3 times to remove OP50 bacteria and 1 ml of TRIzol™ Reagent (Thermo #15596018) was added and kept at −80 °C. After 16-24 hrs freezing, the worms were thawed at room temperature to burst the thick cuticle of *C. elegans*. After freeze-thawing, 0.2ml of chloroform (SRL #96764) was added and mixed by mild shaking and incubated for 3 minutes. After that, the mixture was centrifuged at 14,000g for 16 min at 4°C. 500μL of the aqueous phase was collected and 500 μL isopropanol (SRL #38445) was added to it and incubated for 10 min, followed by centrifugation at 14,000g for 10min at 4°C. RNA pellet was washed with 1 mL of 75% ice-cold ethanol followed by centrifugation at 14,000g for 6min at 4°C and then was dissolved in 25 μL DEPC-treated water. cDNA was prepared from 1 μg of RNA using iScript™ cDNA Synthesis Kit.

Real-time PCR was performed with SYBR green fluorophore (iTaq™ Universal SYBR® Green Supermix) using BioRad CFX-96 Real Time PCR System. Relative transcript level of all the genes was normalized against RPL35 gene. Gene-specific primers used in this study are listed in Table-S2

### Lysotracker staining in *C. elegans*

LysoTracker Red DND 99 (Thermo #L7528) was used at a final concentration of 10 µM for in vivo staining. Synchronized L4 worms were staged on respective conditioned NGM plates and were grown for 5 days at 20 °C. Mixed stage population worms were washed three times with M9 buffer to remove OP50 bacteria, which served as the food source for *C. elegans*; 100 µL of the worm precipitate was mixed with 1 µL LysoTracker (1mM solution) and incubated for 1 h at 20 °C in the dark. Worms were immobilized on 3% agar pads containing 0.5mM sodium azide, and were imaged using a Leica confocal microscope.

## Acknowledgement

We thank all members of the Gupta lab for insightful suggestions. This work was supported by DBT-Wellcome Trust India Alliance Fellowship (IA/I/16/1/502369), STARS-2 Grant (2023-0210) from the Ministry of Education and IISER K intramural funding to A.G. KC is supported by a Dept of Science and Technology grant (DST/TDT/TC/RARE/2022/32).

D.B and B.M. are supported by a pre-doctoral fellowship from the University Grants Commission and Council of Scientific and Industrial Research, respectively, the authors also acknowledge the IISER Kolkata Central Instrumentation Facility (iCIF) for support and instrumental facilities, and IISER Kolkata for the ICP-MS facility.

## Author contribution statement

DB: Conceptualization, data curation, formal analysis

KC: Conceptualization, data curation, formal analysis, Writing-original draft

RP: Data curation

BM: Resources

AB: Resources, Supervision, Validation, Writing-original draft

AG: Conceptualization, Resources, Supervision, Validation, Writing-original draft, Writing-review and editing, Project administration

## Legends to supplementary figures

**Fig.S1**. **Localization of ATP7A following CuCl_2_+TTM and CdCl_2_+TTM treatment in A549 cells.** (A) Localization of ATP7A (red) with Golgin 97 (green). A549 cells were treated with increasing concentration (20μM, 50μM and 100μM) of CdCl_2_ followed by treatment with 50μM of TTM. Since TTM does not chelate Cd(II), ATP7A retained a similar vesicular localization pattern (B). A549 cells were treated with increasing concentration (20μM, 50μM and 100μM) of CdCl_2_ followed by treatment with 50μM of TTM. Since TTM does not chelate Cd(II), ATP7A retained a similar vesicular localization pattern., Scale bar: 5μm

**Fig.S2**. **Localization of ATP7A following CuCl_2_ and CdCl_2_ treatment in HEK293 cells.**(A) Localization of ATP7A (red) with Golgin-97 (green) under basal conditions (upper panel). ATP7A predominantly colocalizes with the Golgin-97 marker. Upon CuCl_2_ treatment, ATP7A undergoes vesicularization (middle panel). Following CuCl_2_ treatment and subsequent treatment with TTM, ATP7A traffics back to the TGN (lower panel).(B) HEK293 cells were treated with 20μM, 50μM, and 100μM CdCl_2_. ATP7A similarly undergoes vesicularization in a dose-dependent manner. (C) HEK293 were treated with 20μM, 50μM, and 100μM CdCl_2_ followed by treatment with 50μM TTM. Since TTM does not chelate Cd(II), ATP7A retained a similar vesicular localization pattern in both the presence and absence of TTM treatment Scale bar: 5μm

**Fig.S3. Localization of ATP7A following CuCl_2_ and CdCl_2_ treatment in HeLa cells.**(A) Localization of ATP7A (red) with Golgin-97 (green) under basal conditions (upper panel). ATP7A predominantly colocalizes with the Golgin-97 marker. Upon CuCl_2_ treatment, ATP7A undergoes vesicularization (middle panel). Following CuCl_2_ treatment and subsequent treatment with TTM, ATP7A traffics back to the TGN (lower panel).(B) HeLa cells were treated with 20μM, 50μM, and 100μM CdCl_2_. ATP7A similarly undergoes vesicularization in a dose-dependent manner. (C) HeLa cells were treated with 20μM, 50μM, and 100μM CdCl_2_ followed by treatment with 50μM TTM. Since TTM does not chelate Cd(II), ATP7A retained a similar vesicular localization pattern in both the presence and absence of TTM treatment Scale bar: 5μm

**Fig.S4. Localization of ATP7A following ZnCl_2_, CaCl_2_, and MgCl_2_ treatment in A549 cells.** A549 cells are treated with 50μM of ZnCl_2_ (upper panel), 50μM of CaCl_2_ (middle panel) and 50μM of MgCl_2_ (lower panel).In each condition ATP7A (red) is colocalizing with TGN marker Golgin-97 (green). Scale bar-5μm

**Fig.S5. ATP7A does not localize to EEA1- or VPS35-positive compartments under basal conditions or following CuCl_2_ or CdCl_2_ treatment in A549 cells.** (A) Localization of ATP7A (green) with EEA1 (red) under basal conditions (upper panel). ATP7A does not colocalize with the EEA1 (early endosome marker). Upon treatment with 25μM (middle panel) and 50μM (lower panel) CuCl_2_, ATP7A does not show colocalization with EEA1.(B) Similarly, A549 cells were treated with 20μM (upper panel) and 50μM (lower panel) CdCl_2_. ATP7A doesn’t show colocalization with EEA1 under these conditions (C) Localization of ATP7A (green) with VPS35 (red) under basal conditions (upper panel). ATP7A does not colocalize with the VPS35 (retromer positive endosome). Upon treatment with 25μM (middle panel) and 50μM (lower panel) CuCl_2_, ATP7A does not show colocalization with VPS35.(D) Similarly, A549 cells were treated with 20μM (upper panel) and 50μM (lower panel) CdCl_2_. ATP7A did not show colocalization with VPS35 under these conditions. Scale bar: 5μm

**Fig.S6. ATP7A does not localize to RAB 7- or AP-3 positive vesicles under basal conditions or following CuCl_2_ or CdCl_2_ treatment in A549 cells.** (A) Localization of ATP7A (green) with RAB7 (red) under basal conditions (upper panel). ATP7A does not colocalize with the RAB 7 (late endosome marker). Upon treatment with 25μM (middle panel) and 50μM (lower panel) CuCl_2_, ATP7A does not show colocalization with RAB7.(B) Similarly, A549 cells were treated with 20μM (upper panel) and 50μM (lower panel) CdCl_2_. ATP7A did not show colocalization with RAB 7 under these conditions (C) Localization of ATP7A (green) with AP-3 (red) under basal conditions (upper panel). ATP7A does not colocalize with the AP-3 (retromer positive endosome) positive vesicles. Upon treatment with 25 μM (middle panel) and 50μM (lower panel) CuCl_2_, ATP7A does not show colocalization with AP-3.(D) Similarly, A549 cells were treated with 20μM (upper panel) and 50μM (lower panel) CdCl_2_. ATP7A doesn’t show colocalization with AP-3 under these conditions. Scale bar: 5μm

**Fig.S7. Localization of ATP7A with RAB11 following CuCl_2_ and CdCl_2_ treatment in A549 cells.** (A) Localization of ATP7A (green) with RAB11 (red) under basal conditions (upper panel). ATP7A shows little perinuclear colocalization with the RAB11 (late recycling endosome marker) vesicles. Upon treatment with 25μM (middle panel) and 50μM (lower panel) CuCl_2_, ATP7A shows decreased colocalization with RAB11 in a dose dependent manner. (B) Similarly, A549 cells were treated with 20μM (upper panel) and 50μM (lower panel) CdCl_2_. ATP7A shows decreased colocalization with RAB11 under these conditions in a dose dependent manner (C) PCC values are quantified from 23 cells (basal), 27 cells (25μM CuCl_2_-treated),22 cells (50μM CuCl_2_-treated), 27 cells (25 μM CdCl_2_ treated), 33 cells (50μM CdCl_2_ treated). p values for each condition are indicated in the graph. Scale bar: 5μm

**Fig.S8. Level of Cadmium accumulation in A549 cells treated with increasing concentrations of CdCl_2_ for 2 hours-** A549 cells are treated with 20μM, 50μM, and 100μM CdCl_2_ for 2 hours. Intracellular Cd levels were measured by ICP-MS and represented as ng Cd/1.5 × 10^5 cells. Under basal conditions, the Cd level is 0.060 ng/1.5 × 10^5 cells. Upon treatment with 20μM CdCl₂, intracellular Cd accumulation increased to 5.25 ng/1.5 × 10^5 cells, while treatment with 50μM and 100μM CdCl_2_ resulted in Cd accumulation levels of 8.45275 ng/1.5 × 10^5 cells and 14.10975 ng/1.5 × 10^5 cells, respectively. Number of biological replicates = 2; number of technical replicates = 5 for each condition. p values for each condition are written on the graph.

**Fig.S9**. **Localization of ATP7B following ZnCl_2_, CaCl_2_,and MgCl_2_ treatment in HepG2 cells** HepG2 cells are treated with 50μM of ZnCl_2_ (upper panel), 50μM of CaCl_2_ (middle panel) and 50μM of MgCl_2_ (lower panel).In each condition ATP7B (green) is colocalizing with TGN marker p230 (red). Scale bar: 5μm

**Fig.S10. Localization of ATP7B with EEA1 following CuCl_2_ and CdCl_2_ treatment in HepG2 cells.** (A) Localization of ATP7B (green) with EEA1 (red) under basal conditions (upper panel). ATP7B doesn’t colocalize with the EEA marker. Upon treatment with 50μM (middle panel) and 100μM (lower panel) CuCl_2_, ATP7B does not show colocalization with EEA1.(B) Similarly, HepG2 cells were treated with 20μM, 50μM and 100μM CdCl_2_. No colocalization with EEA1 has been observed. Scale bar: 5μm

