## Supplemental file 1 for "Copper-transporting ATPase ATP7B and the lysosomal exocytosis pathway synergise to detoxify cadmium from cells"

**Legends to supplementary figures:**

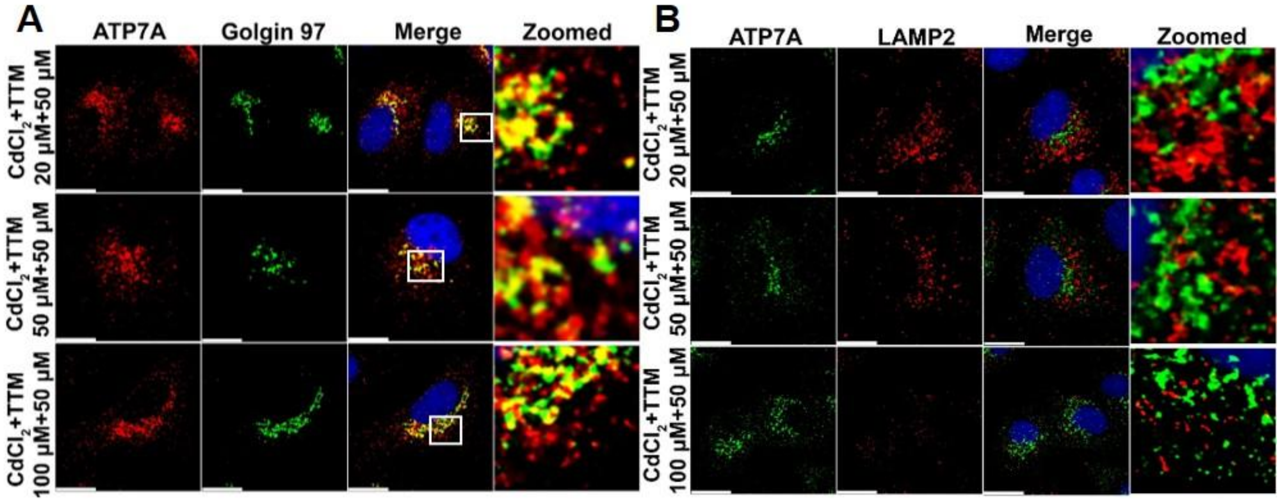


**Fig.S1. Localization of ATP7A following $CuCl_2$+TTM and $CdCl_2$+TTM treatment in A549 cells.** (A) Localization of ATP7A (red) with Golgin 97 (green). A549 cells were treated with increasing concentration (20µM, 50µM and 100µM) of $CdCl_2$ followed by treatment with 50µM of TTM. Since TTM does not chelate Cd(II), ATP7A retained a similar vesicular localization pattern (B). A549 cells were treated with increasing concentration (20µM, 50µM and 100µM) of $CdCl_2$ followed by treatment with 50µM of TTM. Since TTM does not chelate Cd(II), ATP7A retained a similar vesicular localization pattern., Scale bar: 5µm
