## Supplemental file 2 for "Copper-transporting ATPase ATP7B and the lysosomal exocytosis pathway synergise to detoxify cadmium from cells"

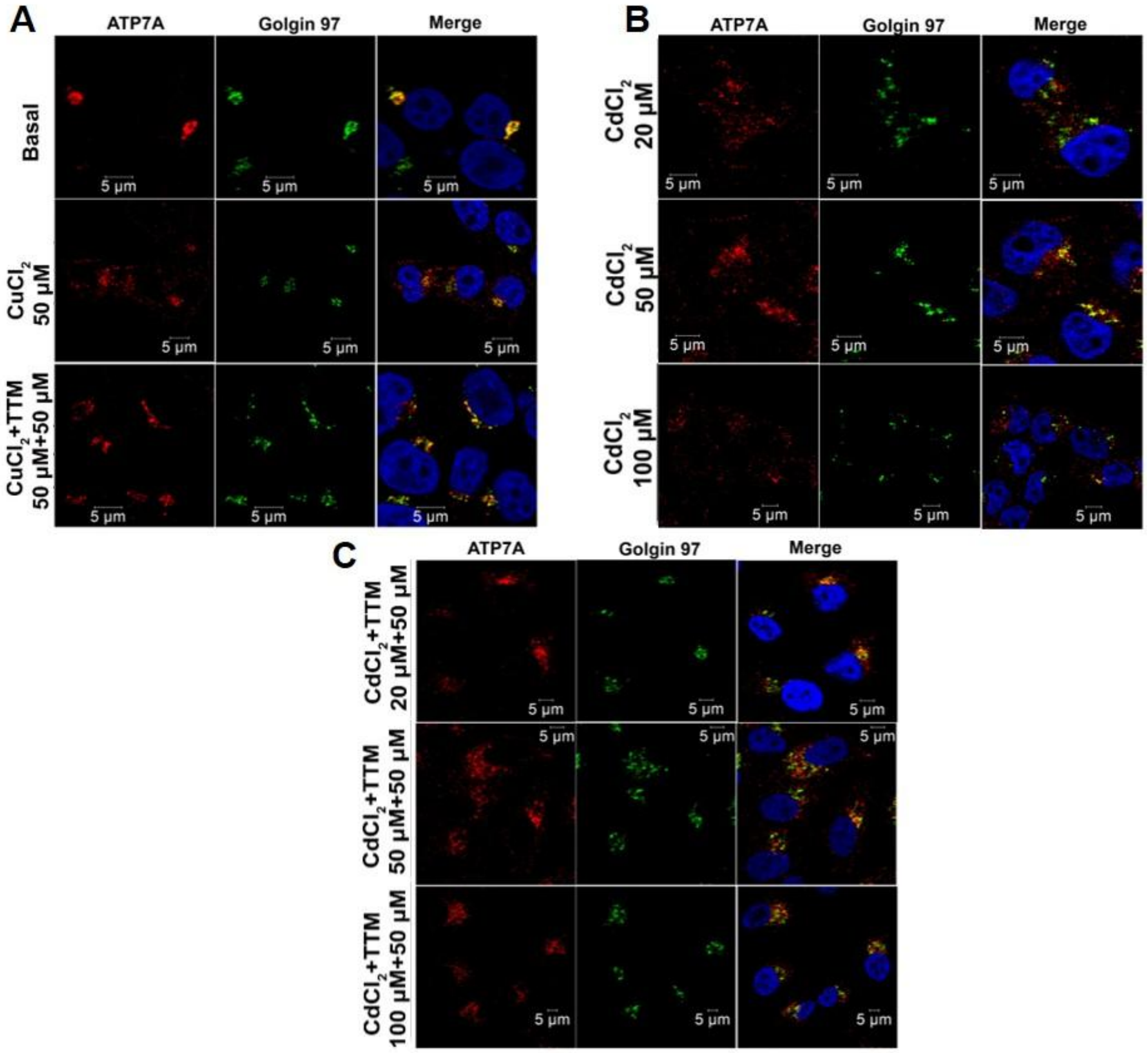


**Fig.S2**. **Localization of ATP7A following $CuCl_2$ and $CdCl_2$ treatment in HEK293 cells.**(A) Localization of ATP7A (red) with Golgin-97 (green) under basal conditions (upper panel). ATP7A predominantly colocalizes with the Golgin-97 marker. Upon $CuCl_2$ treatment, ATP7A undergoes vesicularization (middle panel). Following $CuCl_2$ treatment and subsequent treatment with TTM, ATP7A traffics back to the TGN (lower panel).(B) HEK293 cells were treated with 20µM, 50µM, and 100µM $CdCl_2$. ATP7A similarly undergoes vesicularization in a dose-dependent manner. (C) HEK293 were treated with 20µM, 50µM, and 100µM $CdCl_2$ followed by treatment with 50µM TTM. Since TTM does not chelate Cd(II), ATP7A retained a similar vesicular localization pattern in both the presence and absence of TTM treatment Scale bar: 5µm
