## Supplemental file 3 for "Copper-transporting ATPase ATP7B and the lysosomal exocytosis pathway synergise to detoxify cadmium from cells"

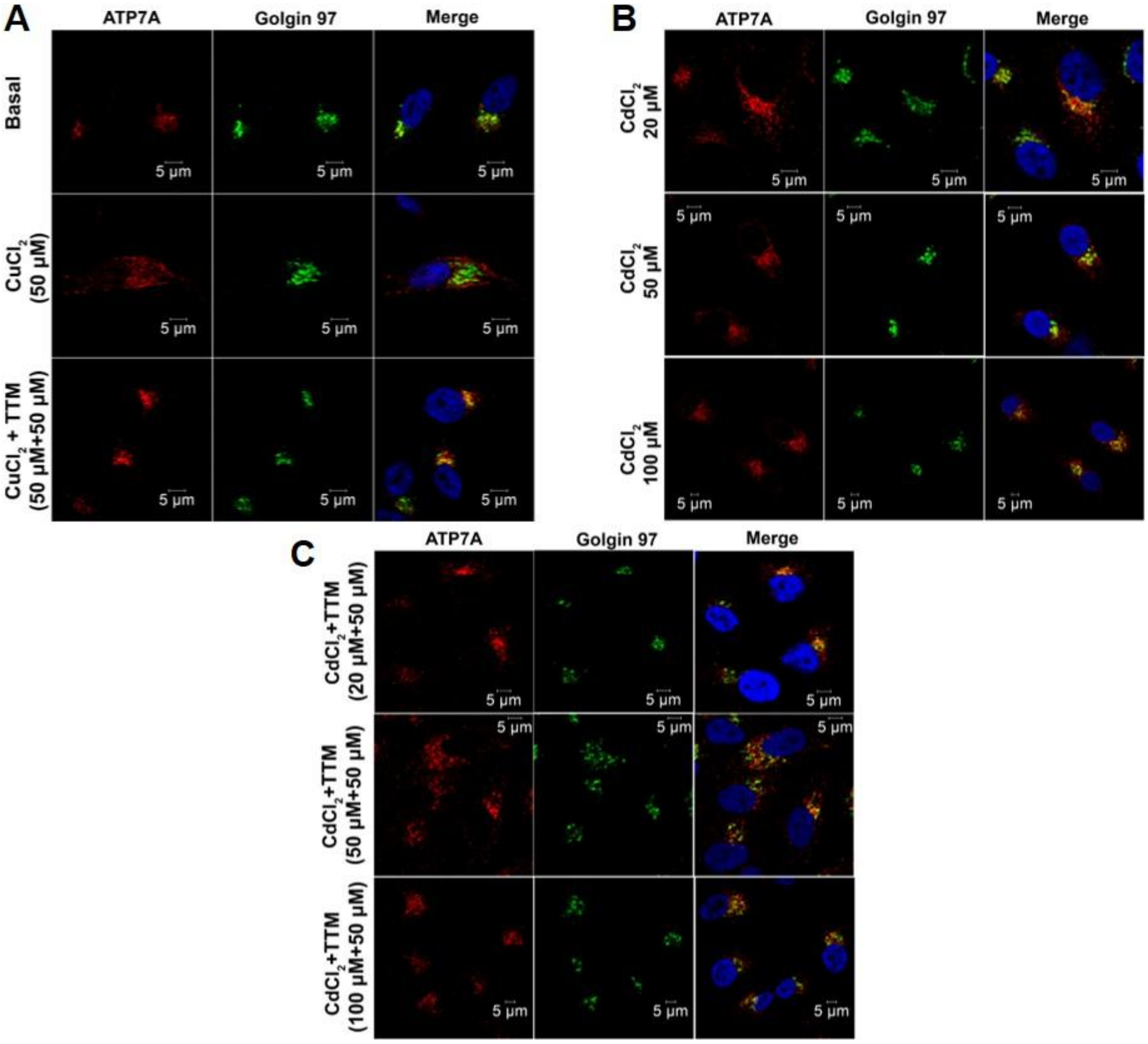


**Fig.S3. Localization of ATP7A following $CuCl_2$ and $CdCl_2$ treatment in HeLa cells.**(A) Localization of ATP7A (red) with Golgin-97 (green) under basal conditions (upper panel). ATP7A predominantly colocalizes with the Golgin-97 marker. Upon $CuCl_2$ treatment, ATP7A undergoes vesicularization (middle panel). Following $CuCl_2$ treatment and subsequent treatment with TTM, ATP7A traffics back to the TGN (lower panel).(B) HeLa cells were treated with 20μM, 50μM, and 100μM $CdCl_2$. ATP7A similarly undergoes vesicularization in a dose-dependent manner. (C) HeLa cells were treated with 20μM, 50μM, and 100μM $CdCl_2$ followed by treatment with 50μM TTM. Since TTM does not chelate Cd(II), ATP7A retained a similar vesicular localization pattern in both the presence and absence of TTM treatment Scale bar: 5μm
