## Supplemental file 4 for "Copper-transporting ATPase ATP7B and the lysosomal exocytosis pathway synergise to detoxify cadmium from cells"

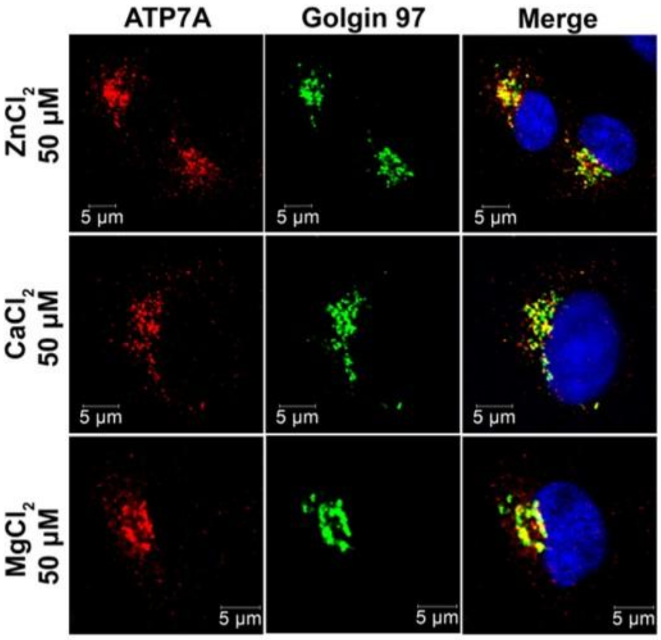


**Fig.S4. Localization of ATP7A following $ZnCl_2$, $CaCl_2$, and $MgCl_2$ treatment in A549 cells.** A549 cells are treated with 50μM of $ZnCl_2$ (upper panel), 50μM of $CaCl_2$ (middle panel) and 50μM of $MgCl_2$ (lower panel).In each condition ATP7A (red) is colocalizing with TGN marker Golgin-97 (green) .Scale bar-5μm
