## Supplemental file 5 for "Copper-transporting ATPase ATP7B and the lysosomal exocytosis pathway synergise to detoxify cadmium from cells"

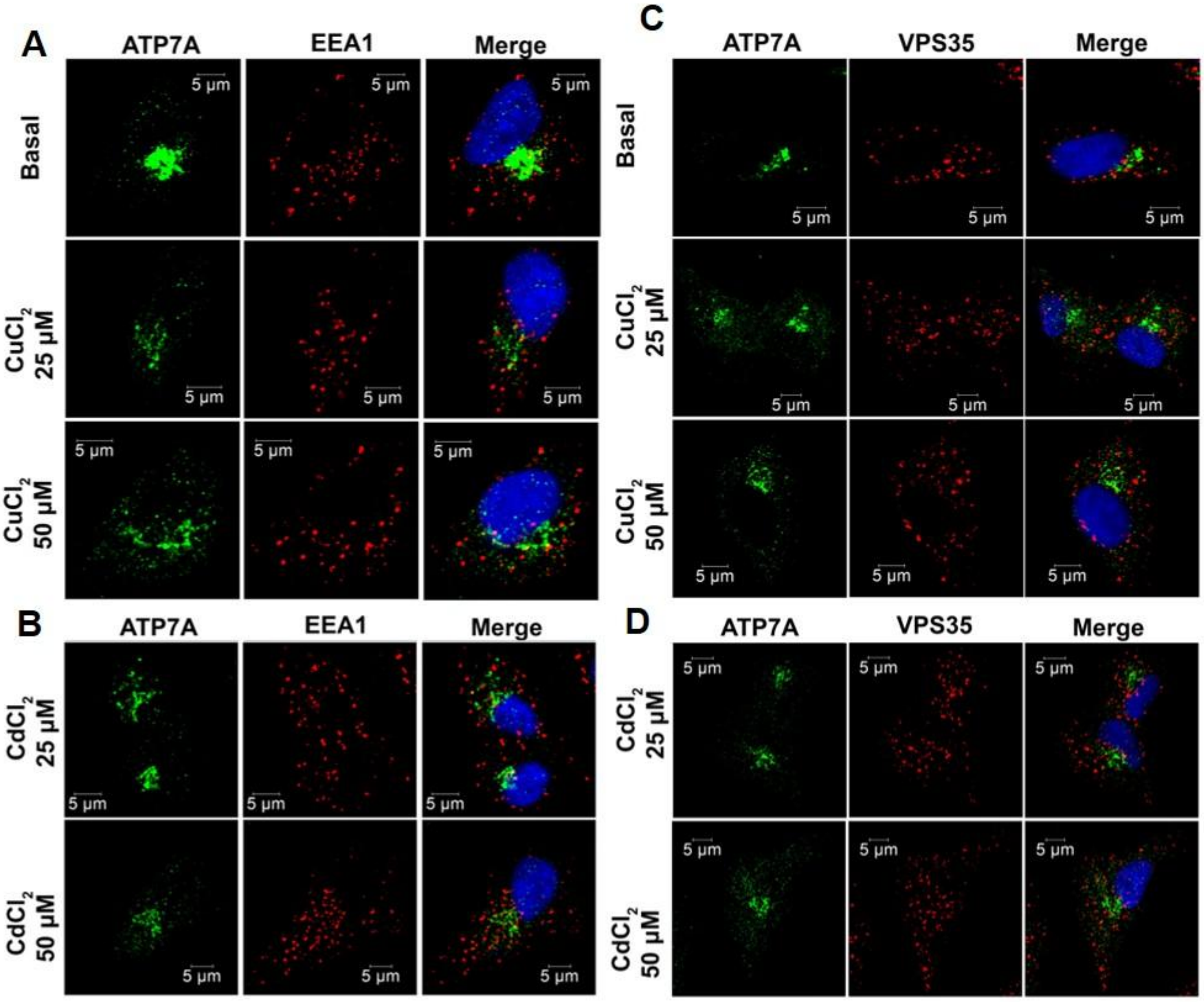


**Fig.S5. ATP7A does not localize to EEA1- or VPS35-positive compartments under basal conditions or following $CuCl_2$ or $CdCl_2$ treatment in A549 cells.** (A) Localization of ATP7A (green) with EEA1 (red) under basal conditions (upper panel). ATP7A does not colocalize with the EEA1 (early endosome marker). Upon treatment with 25μM (middle panel) and 50μM (lower panel) $CuCl_2$, ATP7A does not show colocalization with EEA1.(B) Similarly, A549 cells were treated with 20μM (upper panel) and 50μM (lower panel) $CdCl_2$. ATP7A doesn't show colocalization with EEA1 under these conditions (C) Localization of ATP7A (green) with VPS35 (red) under basal conditions (upper panel). ATP7A does not colocalize with the VPS35 (retromer positive endosome). Upon treatment with 25μM (middle panel) and 50μM (lower panel) $CuCl_2$, ATP7A does not show colocalization with VPS35.(D) Similarly, A549 cells were treated with 20μM (upper panel) and 50μM (lower panel) $CdCl_2$. ATP7A did not show colocalization with VPS35 under these conditions. Scale bar: 5μm
