## Supplemental file 6 for "Copper-transporting ATPase ATP7B and the lysosomal exocytosis pathway synergise to detoxify cadmium from cells"

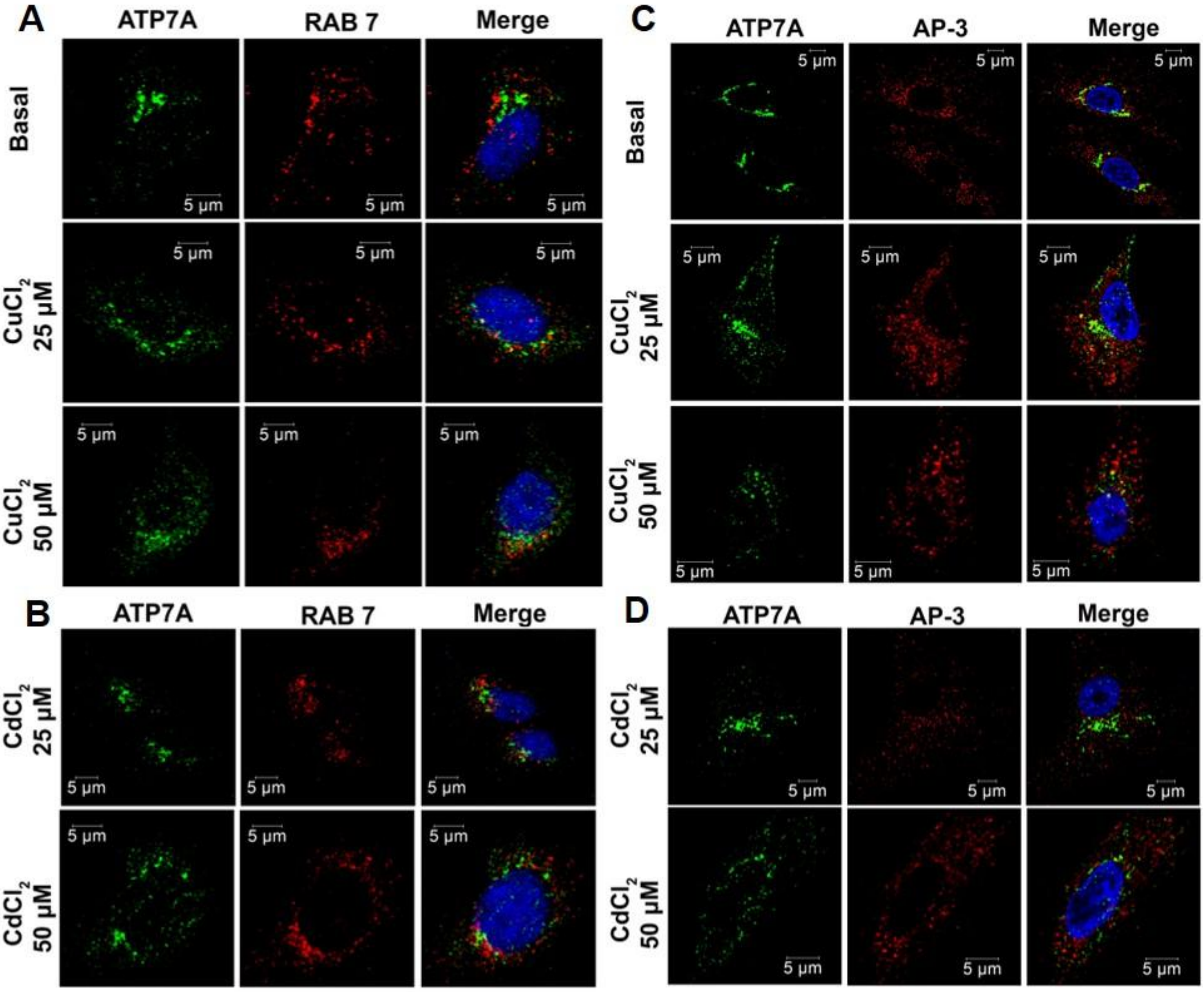


**Fig.S6. ATP7A does not localize to RAB 7- or AP-3 positive vesicles under basal conditions or following $CuCl_2$ or $CdCl_2$ treatment in A549 cells.** (A) Localization of ATP7A (green) with RAB7 (red) under basal conditions (upper panel). ATP7A does not colocalize with the RAB 7 (late endosome marker). Upon treatment with 25μM (middle panel) and 50μM (lower panel) $CuCl_2$, ATP7A does not show colocalization with RAB7.(B) Similarly, A549 cells were treated with 20μM (upper panel) and 50μM (lower panel) $CdCl_2$. ATP7A did not show colocalization with RAB 7 under these conditions (C) Localization of ATP7A (green) with AP-3 (red) under basal conditions (upper panel). ATP7A does not colocalize with the AP-3 (retromer positive endosome) positive vesicles. Upon treatment with 25 μM (middle panel) and 50μM (lower panel) $CuCl_2$, ATP7A does not show colocalization with AP-3.(D) Similarly, A549 cells were treated with 20μM (upper panel) and 50μM (lower panel) $CdCl_2$. ATP7A doesn't show colocalization with AP-3 under these conditions. Scale bar: 5μm
