## Supplemental file 7 for "Copper-transporting ATPase ATP7B and the lysosomal exocytosis pathway synergise to detoxify cadmium from cells"

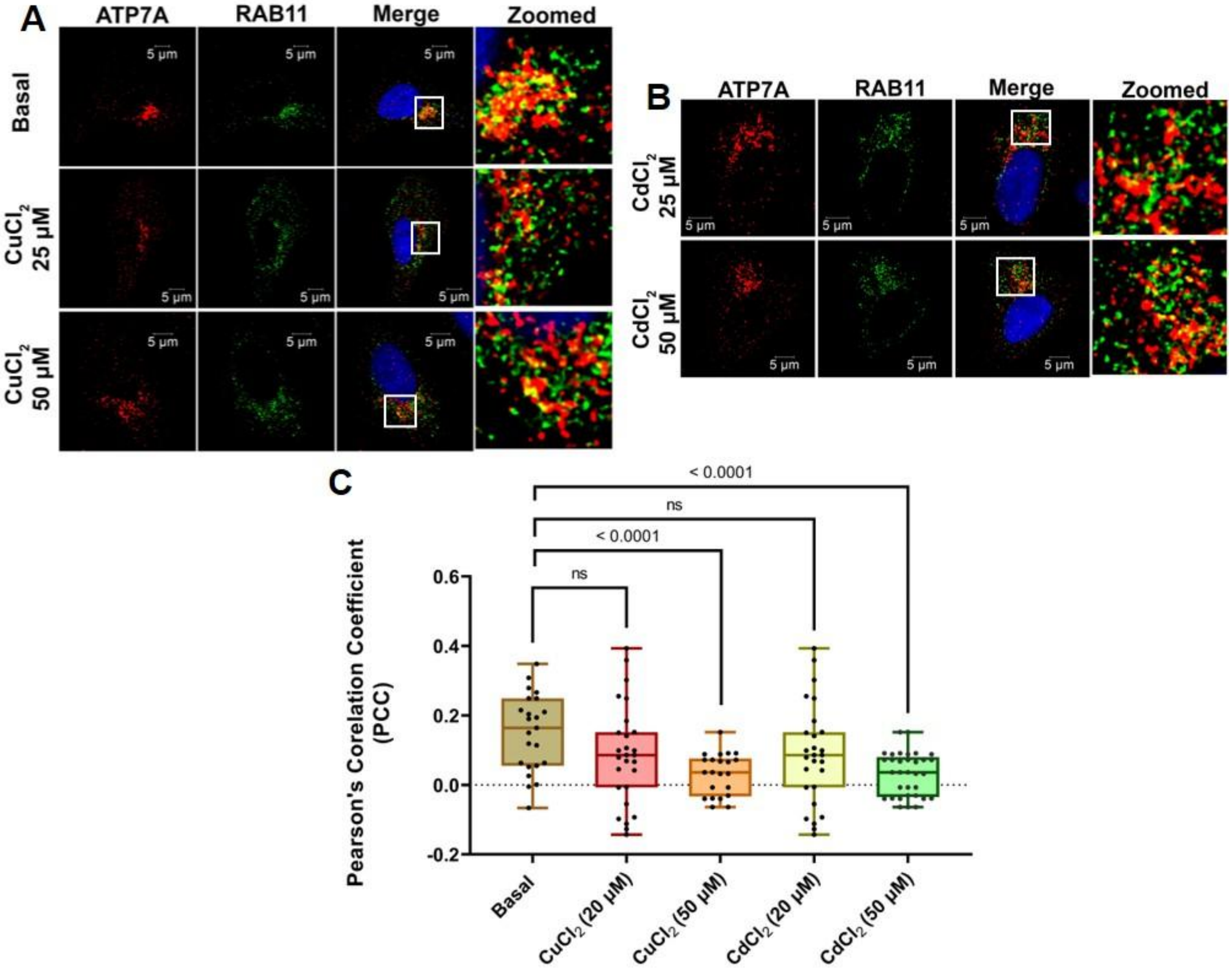


**Fig.S7. Localization of ATP7A with RAB11 following $CuCl_2$ and $CdCl_2$ treatment in A549 cells.** (A) Localization of ATP7A (green) with RAB11 (red) under basal conditions (upper panel). ATP7A shows little perinuclear colocalization with the RAB11 (late recycling endosome marker) vesicles. Upon treatment with 25µM (middle panel) and 50µM (lower panel) $CuCl_2$, ATP7A shows decreased colocalization with RAB11 in a dose dependent manner. (B) Similarly, A549 cells were treated with 20µM (upper panel) and 50µM (lower panel) $CdCl_2$. ATP7A shows decreased colocalization with RAB11 under these conditions in a dose dependent manner (C) PCC values are quantified from 23 cells (basal), 27 cells (25µM $CuCl_2$-treated),22 cells (50µM $CuCl_2$-treated), 27 cells (25 µM $CdCl_2$ treated), 33 cells (50µM $CdCl_2$ treated). p values for each condition are indicated in the graph. Scale bar: 5µm
