## Supplemental file 8 for "Copper-transporting ATPase ATP7B and the lysosomal exocytosis pathway synergise to detoxify cadmium from cells"

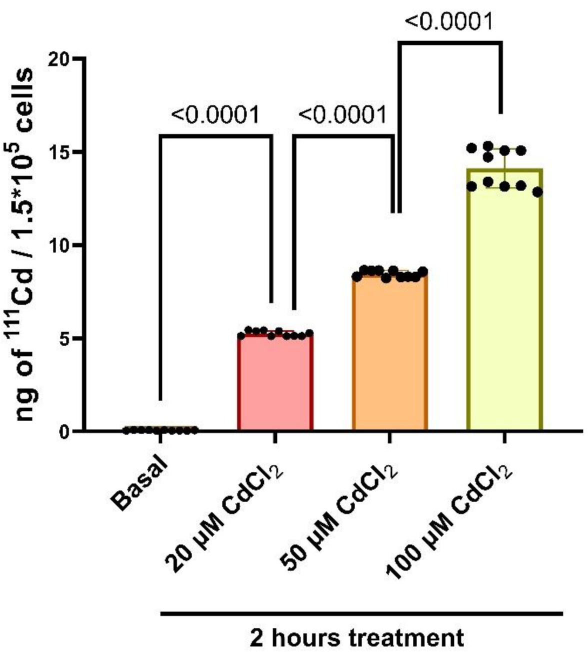


**Fig.S8. Level of Cadmium accumulation in A549 cells treated with increasing concentrations of $CdCl_2$ for 2 hours-** A549 cells are treated with 20µM, 50µM, and 100µM $CdCl_2$ for 2 hours. Intracellular Cd levels were measured by ICP-MS and represented as ng Cd/1.5 × 10^5 cells. Under basal conditions, the Cd level is 0.060 ng/1.5 × 10^5 cells. Upon treatment with 20µM $CdCl_2$, intracellular Cd accumulation increased to 5.25 ng/1.5 × 10^5 cells, while treatment with 50µM and 100µM $CdCl_2$ resulted in Cd accumulation levels of 8.45275 ng/1.5 × 10^5 cells and 14.10975 ng/1.5 × 10^5 cells, respectively. Number of biological replicates = 2; number of technical replicates = 5 for each condition. p values for each condition are written on the graph.
