## Supplemental file 9 for "Copper-transporting ATPase ATP7B and the lysosomal exocytosis pathway synergise to detoxify cadmium from cells"

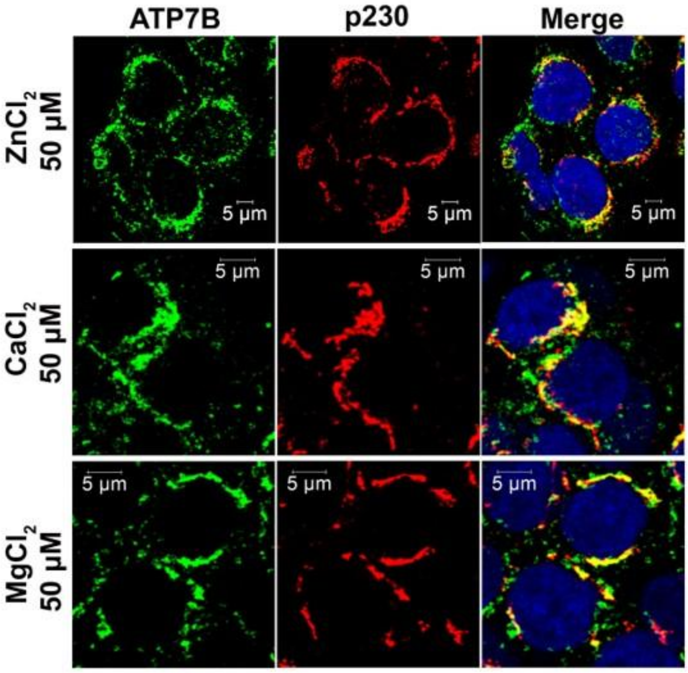


**Fig.S9. Localization of ATP7B following $ZnCl_2$, $CaCl_2$,and $MgCl_2$ treatment in HepG2 cells** HepG2 cells are treated with 50µM of $ZnCl_2$ (upper panel), 50µM of $CaCl_2$ (middle panel) and 50µM of $MgCl_2$ (lower panel).In each condition ATP7B (green) is colocalizing with TGN marker p230 (red). Scale bar: 5µm
