## Supplemental file 10 and Supplemental table 1 and 2 for "Copper-transporting ATPase ATP7B and the lysosomal exocytosis pathway synergise to detoxify cadmium from cells"

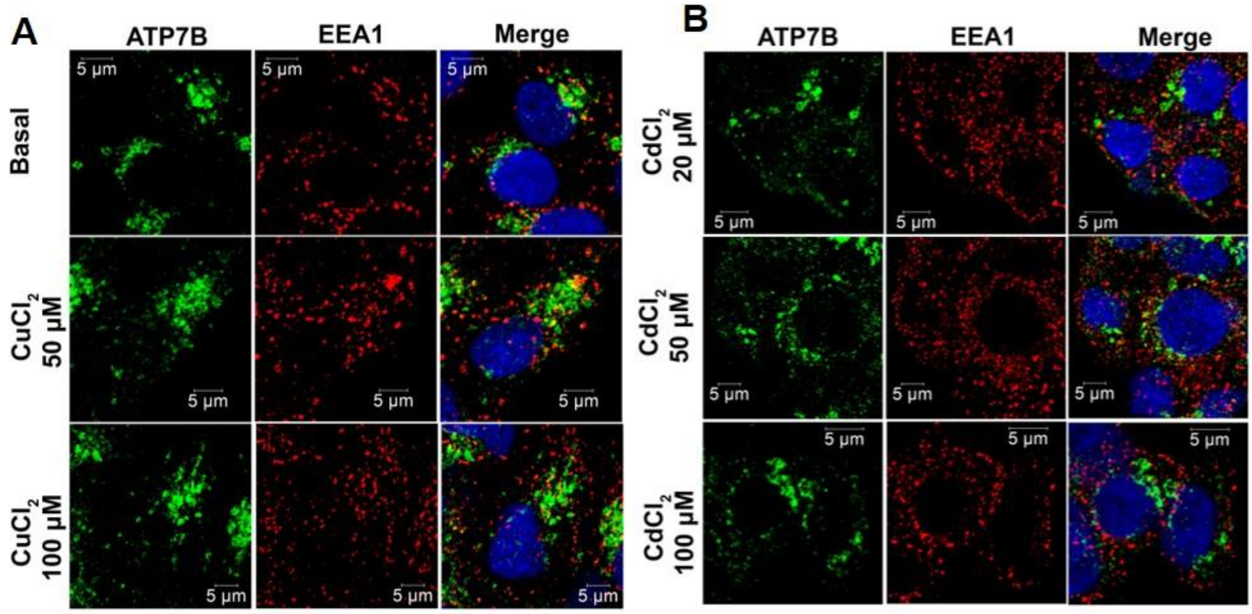


**Fig.S10. Localization of ATP7B with EEA1 following $CuCl_2$ and $CdCl_2$ treatment in HepG2 cells.** (A) Localization of ATP7B (green) with EEA1 (red) under basal conditions (upper panel). ATP7B doesn't colocalize with the EEA marker. Upon treatment with 50µM (middle panel) and 100µM (lower panel) $CuCl_2$, ATP7B does not show colocalization with EEA1.(B) Similarly, HepG2 cells were treated with 20µM , 50µM and 100µM $CdCl_2$. No colocalization with EEA1 has been observed. Scale bar: 5µm

| Table-1 | | |
|---|---|---|
| Name of the Antibody | Make and cat.no. | Dilution used |
| ATP7B | Abcam #ab124973 | 1:400 |
| ATP7A | Santa Cruz#sc376467 | 1:400 |
| LAMP2 | DSHB #H4B4, | 1:200 |
| Golgin 97 | CST#13192S | 1:400 |
| p230 | BD Biosciences #611280 | 1:800 |
| EEA1 | BD Biosciences #610457 | 1:500 |
| RAB11 | Invitrogen # 715300 | 1:300 |
| RAB 7 (for IF) | Santa Cruz#sc271608 | 1:200 |
| RAB 7 (for WB) | Invitrogen# PA5-52369 | 1:1000 |
| Cathepsin D | BD Biosciences # C47620 | 1:1000 |
| VPS35 | Santa Cruz#sc374372 | 1:500 |
| AP3M1 | Abcam# ab201227 | 1:300 |
| TFEB | CST #4240 | 1:200 |
| Goat anti-Rabbit Alexa488 | Thermo #A-11034, | 1:2500 |
| Donkey anti-Mouse Alexa 555 | Thermo #A-11034 | 1:2500 |
| Goat anti-Rabbit Alexa568 | Thermo #A-11011 | 1:2500 |
| α-tubulin | Affinity Biosciences #AF7010 | 1:10000 |
| anti-rabbit-IgG conjugated to HRP | CST#7074 | 1:6000 |
| anti-mouse-IgG conjugated to HRP | CST #7076 | 1:6000 |

| Table-2 | |
|---|---|
| Gene Name | Sequence (5' to 3') |
| *cua-1* | Forward- TGTGAGTCTTCTCGACGGTT |
| | Reverse- GGAGTTGCACGTCATTCCTTTAAT |
| *glo-1* | Forward -TGGTGATCCAGGTGTCGGTA |
| | Reverse -CCATATCGGTCTTGGCCTGA |
| *glo-3* | Forward -GCCAATGACGATCGCAAG |
| | Reverse- GAATGATTTTGGGCTGTATG |
| *apb-3* | Forward- CAACTCTCTCAGATCGCCCG |
| | Reverse -TGGCGTAAGTTCCTTCCTCT |
| *K09C4.5* | Forward- AGAAGTGGCGTCAAGGACAG |
| | Reverse- AAGCAAGCAAACATCGCAGG |
| *pgp-2* | Forward- AGTCAGACAAGTCTAAGCTCTCA |
| | Reverse- ACGGAAAACCAGCTCCATGA |
| *RPL35* | Forward- GCAAGAACATCGCCAGACTCTTG |
| | Reverse- CTTAGCCTGTTGCTTGGCAGATC |
